# Dimeric Cin8 motors have an inherent plus-end bias and weak inter-head coordination

**DOI:** 10.1101/2025.09.24.678381

**Authors:** Himanshu Pandey, Tzu-Chen Ma, Eric Bonventre, Luke M. Rice, Larisa Gheber, William O. Hancock

## Abstract

Kinesin-5 motors are bipolar tetramers that crosslink and slide antiparallel microtubules during mitotic spindle assembly. Fungal kinesin-5 motors, such as Cin8, exhibit bidirectional motility, switching between minus- and plus-end-directed stepping in response to environmental conditions; however, the molecular basis of this directional switching remains unclear. To better understand the origin of this bidirectional behavior, we investigated the motility and ATPase kinetics of two Cin8 dimers, created by fusing the motor domains to a stable coiled-coil domain from kinesin-1. To investigate the role of the proximal neck coiled-coil region in coordinating motor activity, we compared Cin8 dimers that included or lacked the first four heptads of the Cin8 neck-coil domain. By analyzing the stepping kinetics, microtubule residence times, and directional switching dynamics, we found that these Cin8 dimers move processively with a net plus-end directionality along with undirected movements, behaviors that mimic the plus-ended motility state of wild-type Cin8. However, fast minus-ended motility seen in wild-type Cin8 tetramers was not observed in the dimers. The instantaneous velocity distributions and ATPase rates were inconsistent with the undirected movement being solely due to passive diffusion, suggesting that they reflect random bidirectional stepping. Fewer undirected movements were seen on yeast microtubules, their native physiological substrate, compared to on bovine microtubules. Replacing the Cin8 neck-coil domain with a stable coiled-coil led to faster plus-end stepping, fewer undirected movements, a reduction in the microtubule binding duration, and enhanced coupling between ATP hydrolysis and plus-end stepping. Our results suggest that the native Cin8 neck coil confers flexibility between the two motor domains that contributes to bidirectional stepping, and that sustained minus-end movement requires regions outside the motor domain.

**Statement of Significance:** The kinesin-5 family of motors, which contain two pairs of heads located on either end of a long stalk domain, power mitotic spindle formation by sliding antiparallel microtubules. The yeast kinesin-5, Cin8 moves to microtubule minus-ends under specific conditions, breaking the dogma that N-terminal kinesins move to microtubule plus-ends. To gain insight into how Cin8 changes its walking direction, we analyzed engineered Cin8 dimers, models of one half of full-length Cin8. The dimers step erratically, consistent with stepping in both directions, but have a plus-end bias. We find evidence that the two heads poorly coordinate, which contrasts with other kinesins, and suggest that the stepping direction may be regulated by altering the degree of inter-head coordination.

## Introduction

Kinesins are a diverse family of microtubule-based motor proteins that play essential roles in intracellular transport and mitotic spindle dynamics. Canonically, kinesin directionality depends on the position of the motor domain: kinesins with N-terminal motor domains move toward the microtubule plus end, whereas C-terminal kinesins move toward the minus end (1,2). However, this paradigm is challenged by members of the kinesin-5 and kinesin-14 families of mitotic motors that exhibit bidirectional motility (3–7). During elongation of the mitotic spindle, kinesin-5 motors use their two pairs of motor domains to slide apart antiparallel microtubules (8–11), with each pair of heads moving toward microtubule plus-ends (12). However, fungal kinesin-5s, such as Cin8 and Kip1 from *Saccharomyces cerevisiae* and Cut7 from *Schizosaccharomyces pombe*, have also been shown to switch to minus-end directionality under certain conditions (3–5,13). Consistent with this minus-end directionality, Cin8 localizes near the spindle pole bodies at microtubule minus ends in early mitotic cells with monopolar spindles, and this minus-end localization is proposed to enhance microtubule capture from the neighboring spindle pole bodies to set up proper spindle organization for metaphase to proceed (4,13,14). Understanding how bidirectional motility is controlled, particularly in the context of spindle assembly where precise regulation of motor protein activity is critical, remains a major challenge. Furthermore, the lack of a clear mechanism by which kinesins change their stepping direction represents a significant hole in our understanding of kinesin mechanochemistry.

In N-terminal kinesins such as kinesin-1, stepping is driven by ATP-dependent docking of the neck linker domain, a ∼14-18-residue segment at the C-terminal of the motor domain (15). Processive plus-end stepping is thought to be regulated by inter-head tension transmitted through the neck linker domains (16–19). Consistent with this, increasing the length and flexibility of the neck linker domain has been shown to disrupt head-head coordination and reduce processivity (17,20–22). The mechanism by which Cin8 steps to the microtubule minus-end and the role that inter-head coordination plays in minus-end stepping are not known.

A dominant theme that emerges from Cin8 work to date is that perturbations that enhance microtubule affinity bias the motor toward slow plus-end motility, whereas perturbations that reduce the microtubule affinity tend to bias the motor toward fast minus-end directionality. For instance, motor clustering and crosslinking of antiparallel microtubules by Cin8 bias it toward plus-end-directed motility (3,23). In contrast, increasing the ionic strength or deleting the N-terminal extension, loop-8, or the C-terminal tail, all of which can interact with microtubules, bias the motor toward minus-end directionality (3,4,24–26). The underlying mechanism, however, remains unexplained. A recent model proposes that Cin8 inherently steps in both directions, with minus-end stepping dominating at zero load but having a stronger dependence on load such that plus-end stepping dominates under load (23). An alternative model for the related fungal kinesin-5, Cut7, posits that stepping is inherently minus-end directed and crowding on the microtubule surface promotes plus-end stepping by a steric blocking mechanism (14). There is also evidence that directionality of Cin8 is regulated allosterically by regions outside the motor domain such as the tail or the N-terminal non-motor region (24–26).

In the present study, we investigated the role of inter-head coordination in the bidirectional stepping of dimeric Cin8. We hypothesized that directional switching is enabled by loose mechanical coupling between the two motor domains in each Cin8 dimer that comprise the Cin8 tetramer. To test this, we used the stable coiled-coil of kinesin-1 to dimerize Cin8 motor domains with and without their native proximal neck-coil dimerization domain. We compared the stepping speed and directionality on both bovine and yeast microtubules and analyzed the undirected movements to determine the relative contributions of Brownian diffusion and ATP-driven stepping. We find that Cin8 dimers exhibit slow net plus-end motility along with undirected movements. Including the proximal neck-coil of Cin8 leads to a smaller the mean stepping rate, enhanced undirected movements, and increased ATPase rate. These findings are consistent with a model in which loose mechanical coordination between the motor domains, modulated by structural elements of the neck region, plays a critical role in enabling bidirectional motility in fungal kinesin-5.

## Methods

### Protein Constructs

Plasmids to express Cin8 variants were generated using standard PCR and Gibson assembly and verified by DNA sequencing. All constructs are dimers comprising the *S. cerevisiae* Cin8 motor and neck domains (residues 39–530 for Cin8Kin and 39–558 for Cin8_+28NC_Kin), fused to the Drosophila KHC neck-coil and dimerization coiled-coil domain (residues 345–560; Fig. 1A). Constructs were made by inserting the Cin8 motor domains into the K560-AviN plasmid described previously (27). All constructs begin at Cin8 residue 39 and include a C-terminal AviTag™ and His₆ tag for purification.

**Figure 1:**
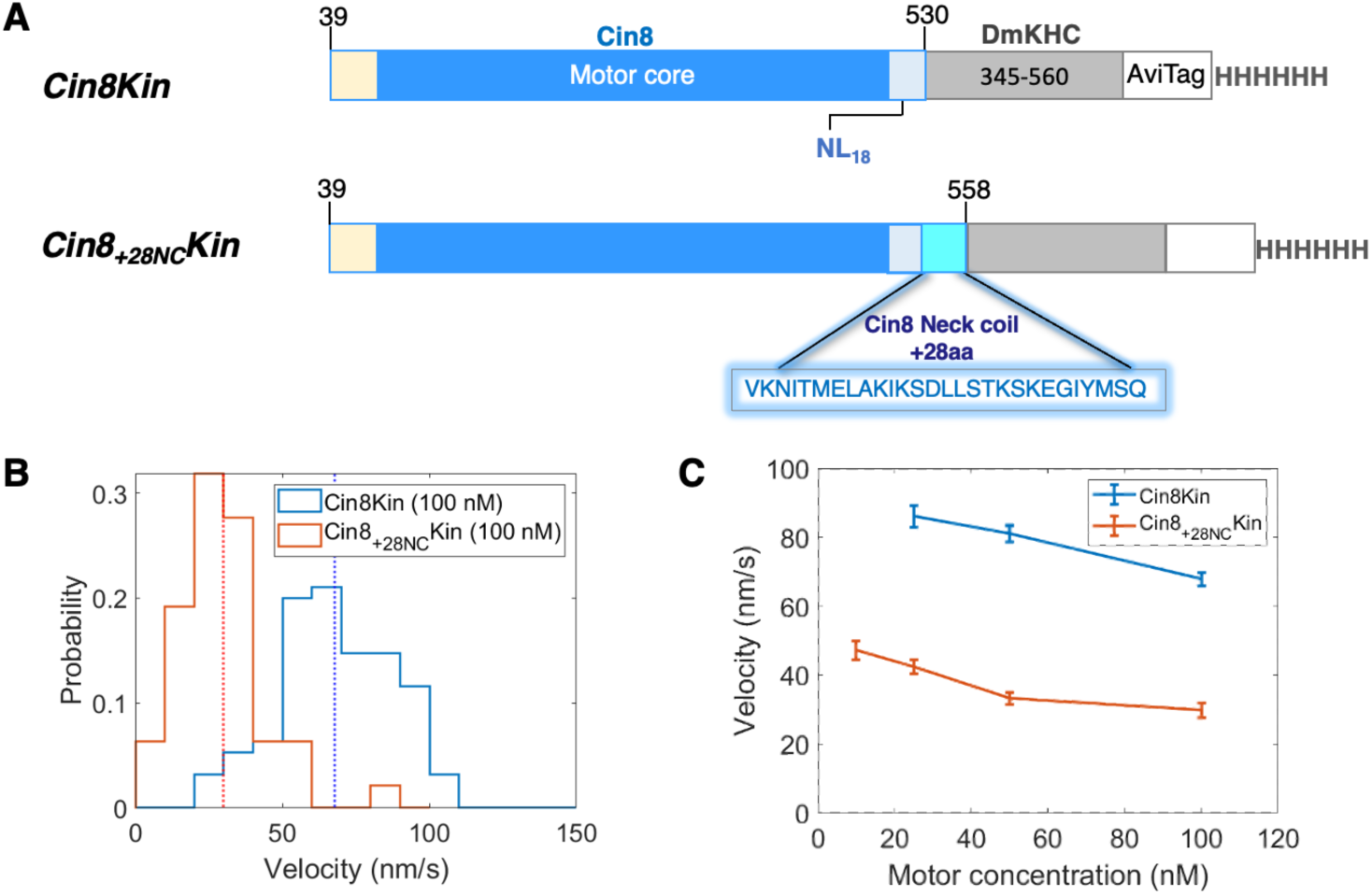
Design and microtubule gliding velocities of dimeric Cin8 constructs. **(A)** Domain organization of Cin8 constructs used in this study. The Cin8 catalytic motor domain (residues 76– 512, blue) is flanked by the N-terminal non-motor region (residues 39–75, yellow) and the C-terminal neck-linker, NL_18_ (residues 513–530, light blue). In Cin8Kin, the Kinesin-1 neck-coil and dimerization domain (residues 345–560, gray) are fused downstream of the Cin8 head, followed by a C-terminal Avi Tag (white) and His_6_ tag. The Cin8_+28NC_Kin construct includes additional 28 residues from the Cin8 neck coil (residues 531–558, cyan), inserted between the Cin8 head and the Kinesin-1 dimerization domain. Diagrams are not to scale **(B)** Distribution of microtubule gliding velocities for both motors with 100 nM motors introduced into flow cell. **(C)** Microtubule gliding velocities at varying motor concentrations in the flow cell. Error bars are SEM with between 47 and 96 microtubules for each motor concentration.

### Purification and Quantification

Bacterial expression procedures were adapted from previously described protocol (25,27). Briefly, constructs were expressed in E. coli BL21-CodonPlus (DE3) grown in 1 L of terrific broth, in-house. 24 mg/l of biotin were added along with the isopropyl 1-thio-β-d-galactopyranoside (IPTG) at the time of induction, followed by nickel gravity column chromatography purification with an 50 mM HEPES based elution buffer containing 2 µM mant-ADP along with 0.25 mM TCEP, 5% glycerol, 500 mM KCl, 10 mM MgCl_2_, 0.01% Triton X-100, pH 7.2. The elution fraction containing motors were supplemented with 10% (w/v) sucrose and 2 mM DTT, flash-frozen and stored at −80 °C. The active motor concentration was determined by measuring the total mant-ADP concentration (356 nm excitation/450 nm emission) in a Shimadzu 3000 spectrofluorometer and correcting for the free mant-ADP, and using a fluorescence enhancement factor of 1.8 for motor-bound mant-ADP (28). Stock active dimeric motor concentrations were in the range of 1–10 μM dimer. All experiments were repeated using motors from at least three independent purification preparations.

Bovine brain tubulin was purified as previously described (29,30). Microtubules were polymerized and stabilized with taxol following standard procedures (29,30). Yeast Tub1 and Tub2 αβ-tubulin was purified from inducibly overexpressing strains of *S. cerevisiae* using Ni-affinity and ion exchange chromatography as previously described (31). Tubulin samples were stored in buffer containing 10 mM PIPES, 1 mM MgCl_2_, 1 mM EGTA, 50 µM GTP, pH 6.9.

### Gliding assay

Flow chambers were assembled and motors adhered to the surface by flowing in 20 μg/ml NeutrAvidin followed by C-terminal biotinylated motors. Taxol-stabilized microtubules were then added to the flow cell in motility buffer (BRB80 adjusted to pH 7.2, 1 mM ATP, 10 μM Taxol, 2 mM MgCl_2_, 5 mM DTT, and 2 mg/ml casein, 0.5 mg/ml BSA). Microtubule movements were monitored with the microscopy setup described for the single-molecule motility assay and recorded at 5 fps. Microtubule polarities were determined by flushing in plus end directed Kif1A in the flow chamber after acquiring the microtubule gliding data on surface immobilized Cin8 dimers. Velocity analysis was limited to only those microtubules that remain within the field during the whole (gliding and Kif1A motility) acquisition. Microtubule velocities were measured by creating kymographs along the MT path using ImageJ-Fiji software and measuring the slope. Periods when microtubules paused or stopped moving were not included in the velocity analysis (Fig. S2).

### Single-molecule motility assay

Motility assays were performed on a Nikon TE2000 TIRF microscope equipped with interference reflection microscopy, at room temperature, as described previously (30,32). Dimeric Cin8 motors were labeled with streptavidin-labeled Quantum dots (Qdot 525; Thermo Fisher) through the C-terminal AviTagTM bound biotin. For labelling, 100 nM dimeric motors were mixed with Qdots at a 1:5 motor:Qdot ratio and incubated in ice for 20 min. An estimated 20% of Avi tags are biotinylated (27); thus, the probability of any Qdot with a motor has more than one motor is <5% (33). Flow cells were functionalized by flowing in 0.5 mg/ml casein, followed by full-length rigor kinesin (27) in the motility buffer (BRB80 adjusted to pH 7.2, 1 mM ATP/ADP, 10 μM Taxol, 2 mM MgCl_2_, 5 mM DTT, and 2 mg/ml casein, 1 mg/ml BSA, 10 mM glucose, 100 μg/ml glucose oxidase, 80 μg/ml catalase 40, 10 mM phosphocreatine, and 50 μg/ml creatine phosphokinase). Taxol-stabilized microtubules, polymerized from bovine or yeast tubulin, were then introduced into the flow cell, where they attached to surface-immobilized rigor kinesin. Subsequently, Qdot labeled motors were introduced and imaged. To accurately measure run times, motors were introduced into the flow cell immediately upon initiating image acquisition, ensuring that no motors were bound prior to the start of imaging. Movies, typically 1000 frames, were recorded at 5 fps. When measuring run durations, motors that landed during the final 20% of the movie were excluded from analysis to prevent censoring by termination of the video. At the end of each experiment, microtubule polarities were determined by flushing in fluorescent KIF1A motors and observing their plus-end motility (28). All microtubules in this study were unlabeled and were imaged by interference reflection microscopy.

### Data analysis

Movies of single Cin8 motors moving along microtubules were analyzed with FIESTA software (34) to generate 1D displacement versus time trajectories. Molecules with no detectable movement were not analyzed and segments of traces where movement ceased for the remaining duration of the movie were excluded as well. Traces were categorized as directed or undirected. Undirected traces were defined as those with overall velocity < 20 nm/s and net displacements < 1 µm (the majority of which were < 0.5 µm). Instantaneous velocities were calculated by fitting a linear regression over a 1-s or 5-s window using the Matlab function, polyfit with n = 1. Velocities were calculated for every point in the timeseries except the first and last points that were limited by the window size. This oversampling approach avoided any randomness due to specifying the precise point of the starting window. For computing the SEM, the number of independent velocity windows was calculated by dividing the entire trace duration by the size of the time window (1 or 5 s). In a control experiment, static particles had average SD of position of 19.8 nm over an entire movie. This translates to a ∼20 nm/s or ∼4 nm/s error for 1-s and 5-s window velocity determinations.

For mean-squared displacement (MSD) analysis, we calculated the squared displacement during each time lag (𝜏) for each tracked particle and averaged the squared displacements for each 𝜏 over all particles. To estimate the diffusion coefficients (D) we fit MSD data with 𝑣^2^𝜏^2^+ 2𝐷𝜏 + 𝑠^2^, where v is mean velocity obtained from the 5 s instantaneous velocity distribution and s set to 10 nm based on the point spread function measurement error in FIESTA. For Gaussian Mixture Model fitting, we used Matlab built-in function “fitgmdist” to fit the PDF distribution of 5-s instantaneous velocities with 3 Gaussian functions. The maximum iteration was set to 1000 to ensure convergence of the fit. All the fitting and plotting were performed using Matlab. Kymographs were generated using ImageJ-FIJI (35).

### Steady state ATPase assay

Cin8 ATPase activity at varying [Mt] was quantified by an enzyme-coupled assay following a protocol adapted from Huang and Hackney (28,36). BRB80 buffer supplemented with 1 mM Mg-ATP, 2 mm phosphoenolpyruvate, 1 mm MgCl_2_, 0.2 mg/ml casein, 10 μM Taxol, 0.25 mM NADH, and 1.5% volume % of Pyruvate Kinase/Lactate dehydrogenase (Sigma P-0294) was used for ATPase assays at 21 °C. Dimeric motor concentrations between 50 and 100 nM were used. NADH absorbance at 340 nm was measured over time on a Molecular Devices FlexStation3 microplate reader, converted to an ATPase rate by linear fit to regions of steady-state absorbance decrease, and divided by the active dimer-motor concentration to give the total hydrolysis cycle rate. k_cat_ and K_M_ (MT) were determined from the fit to a Michaelis-Menten equation of ATP turnover rates at various microtubule concentrations. Mean ± SD values were obtained from five independent assays for each Cin8 variant with motors from at least two different purification preparations. Due to the relative difficulty in expressing and purifying yeast tubulin and the large consumption of microtubules, all ATPase measurements were carried out using bovine brain microtubules.

### Simulations

Cin8 stepping simulations used a Gillespie stochastic stepping model based on an algorithm developed in previous work (37,38). Plus-end directed stepping was the only reaction considered, with the unloaded stepping rate, k^0^_forward_ taken from the mean velocity on yeast microtubules, assuming an 8 nm step size. To model diffusion, a diffusive term was added to each step based on sampling from a normal distribution with mean of zero and standard deviation 𝜎 = √2𝐷𝜏, where D is the chosen diffusion coefficient and 𝜏 is the transition time for that step. All simulations were run 1000 times, and each run was recorded for 50 seconds.

## Results

### Cin8 dimers are plus-end steppers in microtubule gliding assays

To investigate the role of inter-head tension in the directional switching and motility of Cin8, we used a simplified dimeric construct lacking the second pair of motor domains, the tetrameric stalk, and the native tail. To improve solubility, the first 38 amino acids of the unstructured N-terminal domain were deleted. Stable dimers were created by fusing the Cin8 motor domain (amino acids 39–530) to the neck coil and coiled-coil-1 of *Drosophila* kinesin-1 (amino acids 345-560), followed by a biotinylation tag and His_6_tag (Fig. 1A). This dimerization strategy has been used successfully for motors from the kinesin-1, 2, 3, 5, and 7 families and thus allows for direct comparison to previous work (17,28,39–41). The first dimeric construct, Cin8Kin, consists of the Cin8 catalytic core and 18-residue neck linker domain (amino acids 39-530) fused to the kinesin-1 dimerization domain. This construct is similar to one used previously in multimotor gliding assays (25), but lacks N-terminal residues 1-38 and four C-term residues compared to a Cin8Kin construct (residues 1-534) used previously in single-molecule assays (24).

Processive kinesin stepping requires a mechanical connection between the two motor domains, which is achieved through dimerization of the neck-coil domain. In contrast to kinesin-1 (42), the stability of the proximal neck-coil domain of Cin8 is not known. Coiled-coil prediction analyses indicate that the proximal neck coil of Cin8 is less stable than kinesin-1, as evidenced by lower coiled-coil stabilization scores and a reduced number of stabilizing interactions (Fig. S1). Because the inter-head tension that coordinates the hand-over-hand stepping in processive kinesins relies on the proximal neck-coil remaining tightly dimerized, we designed a second construct, Cin8_+28NC_Kin, that included the first four heptads (28-residues) of the native Cin8 neck coil. These constructs were bacterially expressed and purified following standard procedures.

To confirm that the Cin8 dimers are functional, we tested their activity in multi-motor microtubule gliding assays. When biotinylated motors (100 nM solution concentration) were adsorbed to a neutravidin-coated coverslip, Cin8_+28NC_Kin and Cin8Kin steadily drove bovine microtubules in a unidirectional plus-ended manner with no evidence of directional switching. Notably, Cin8Kin exhibited a microtubule gliding velocity approximately twice that of Cin8_+28NC_Kin (67.9 ± 1.9 nm/s versus 29.8 ± 2.1 nm/s; mean ± SEM, N = 95 and 47), meaning that replacing the proximal neck-coil of Cin8 with the stable kinesin-1 neck-coil doubled the apparent stepping rate in the multimotor assay (Fig. 1B). Directionality was confirmed by flowing in GFP-labeled KIF1A motors at the end of the experiment to define the plus end of each microtubule. Previous work using full-length tetrameric Cin8 measured plus-ended gliding velocities ranging from 4.9 nm/s (4) to 57 nm/s (3) and found that microtubule gliding switched to minus-end directionality as the surface motor density was decreased (4). To test this, we decreased the motor concentration in the gliding assay and found no change in directionality (Fig. 1C and S2). Instead, there was a slight increase in plus-end velocity for both motors at reduced motor densities. Thus, in multi-motor assays, Cin8 dimers displayed plus-end directionality, and dimerizing the heads just distal to their neck linker doubled the gliding speed compared to dimers that retained the first four heptads of the native Cin8 neck-coil.

### Single Cin8 dimers move processively with net plus-end directionality

To investigate the speed and directionality of individual Cin8 dimers, we labeled the motors with streptavidin-coated quantum dots and observed their movements along immobilized microtubules polymerized from bovine brain tubulin (Fig. 2A & B). Consistent with the microtubule gliding assays, both Cin8 constructs displayed net plus-end directed movement (Fig. 2A-C & S3), with average trace velocities of 43.8 ± 7.3 nm/s for Cin8_+28NC_Kin and 62.1 ± 6.4 nm/s for Cin8Kin (mean ± SEM; Fig. 2C). However, in contrast to the smooth movements in the multi-motor gliding assay, there were durations of undirected movement, fluctuations in velocity during plus-end directed segments, and frequent short segments with minus-end directionality (Fig. 2A & B). These frequent directional switches across multiple timescales precluded objective classification of processive and diffusive phases and directionality switches. Instead, to comprehensively analyze the single-molecule motility in an unbiased way, we calculated instantaneous velocities for every trace using a 1 s or 5 s window for both motors and plotted the resulting instantaneous velocity distributions (Fig. 2D & E). The 1 s and 5 s results were similar, but due to the long tails in the 1 s distributions we focused our analysis on the 5 s distributions. Both variants had a clear plus-end bias, with Cin8Kin having a distinct peak around ∼100 nm/s and Cin8_+28NC_Kin having a broader distribution that was centered closer to zero. Notably, even with this relatively long 5 s window, 33% of Cin8_+28NC_Kin and 18% of Cin8Kin segments were minus-end directed (Fig. 2E, shaded region), consistent with the considerable undirected movements seen in the raw traces (Fig. 2ACB).

**Figure 2:**
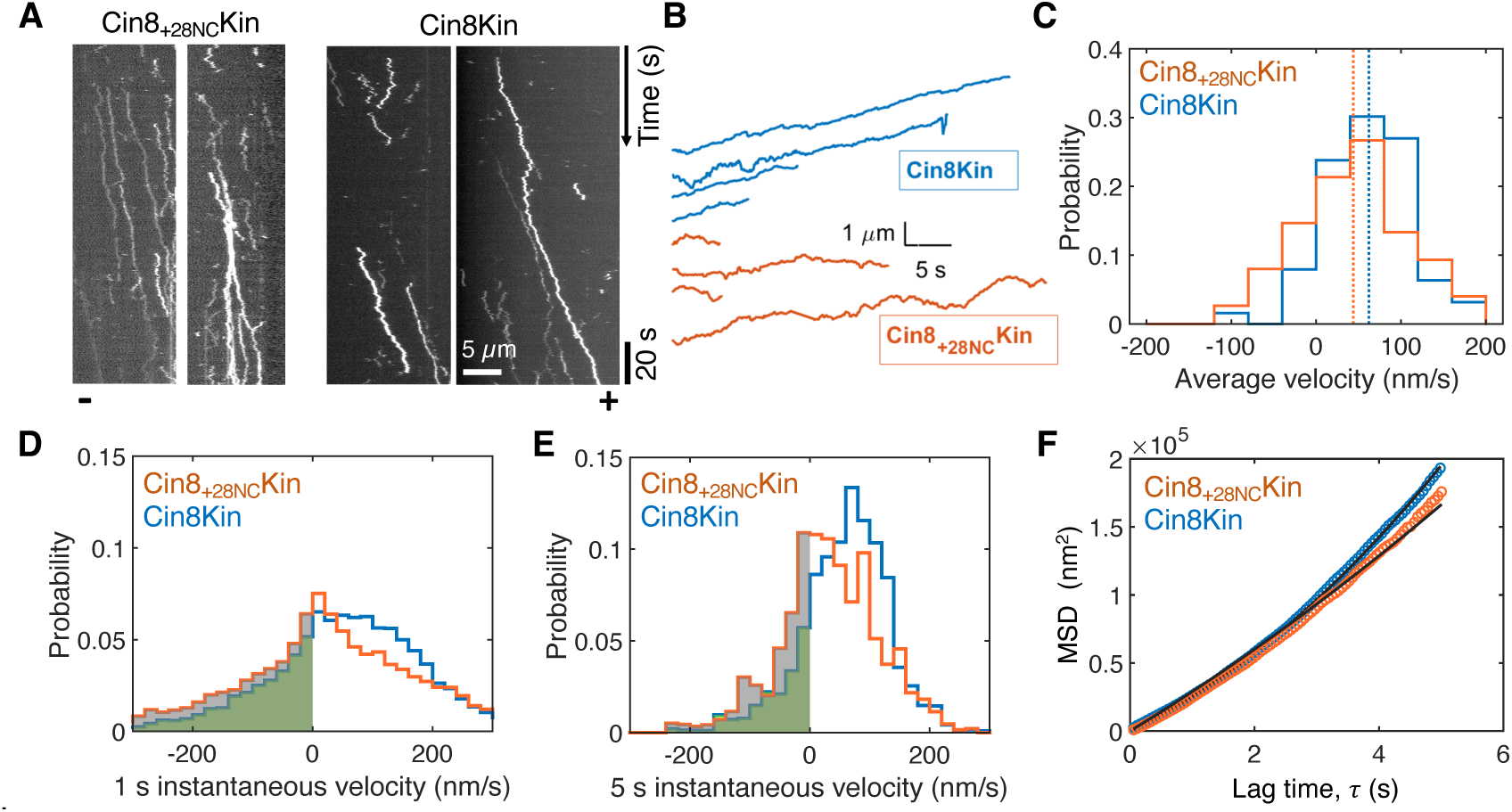
Single-molecule Cin8 dimer motility on bovine microtubules. Representative **(A)** kymographs and **(B)** displacement versus time traces for Cin8_+28NC_Kin (orange) and Cin8Kin (blue) in the presence of 1 mM ATP on surface-immobilized bovine microtubules. Motors were labeled with quantum dots and movies acquired at 5 frames/s. Microtubule polarities, determined by introducing fluorescent KIF1A at the end of the experiment (Fig. S3), are marked at the bottom of kymograph. **(C)** Distribution of whole trace velocities for Cin8_+28NC_Kin and Cin8Kin. Mean velocities were 43.8 ± 7.3 nm/s for Cin8_+28NC_Kin and 62.1 ± 6.4 nm/s for Cin8Kin (mean ± SEM for n= 75 and 63 traces, respectively). **(D)** 1 s and **(E)** 5 s instantaneous velocity distributions for Cin8_+28NC_Kin and Cin8Kin. Average velocities from the 5 s velocity distributions were 32.9 ± 6.2 nm/s for Cin8_+28NC_Kin and 58.6 ± 6.3 nm/s for Cin8Kin (mean ± SEM for N = 516 and 148 independent 5 s windows from 75 and 63 molecules, respectively). Gray (Cin8_+28NC_Kin) and green (Cin8Kin) shaded areas represent minus-end directed velocity segments. **(F)** Mean squared displacement (MSD) analysis in ATP. Data at varying lag time (τ) were fit to MSD = v^2^τ^2^+2Dτ (black lines), with velocities (v) fixed at 32.9 nm/s (Cin8_+28NC_Kin) and 58.6 nm/s (Cin8Kin). Apparent diffusion coefficients (D) were 14338 ± 172 nm²/s and 10921 ± 40 nm²/s, respectively (fit ± 95% CI). Fit to MSD = aτ ^α^ + b gave α of 1.29 for Cin8_+28NC_Kin and 1.38 for Cin8Kin (Table S1).

**Figure 3:**
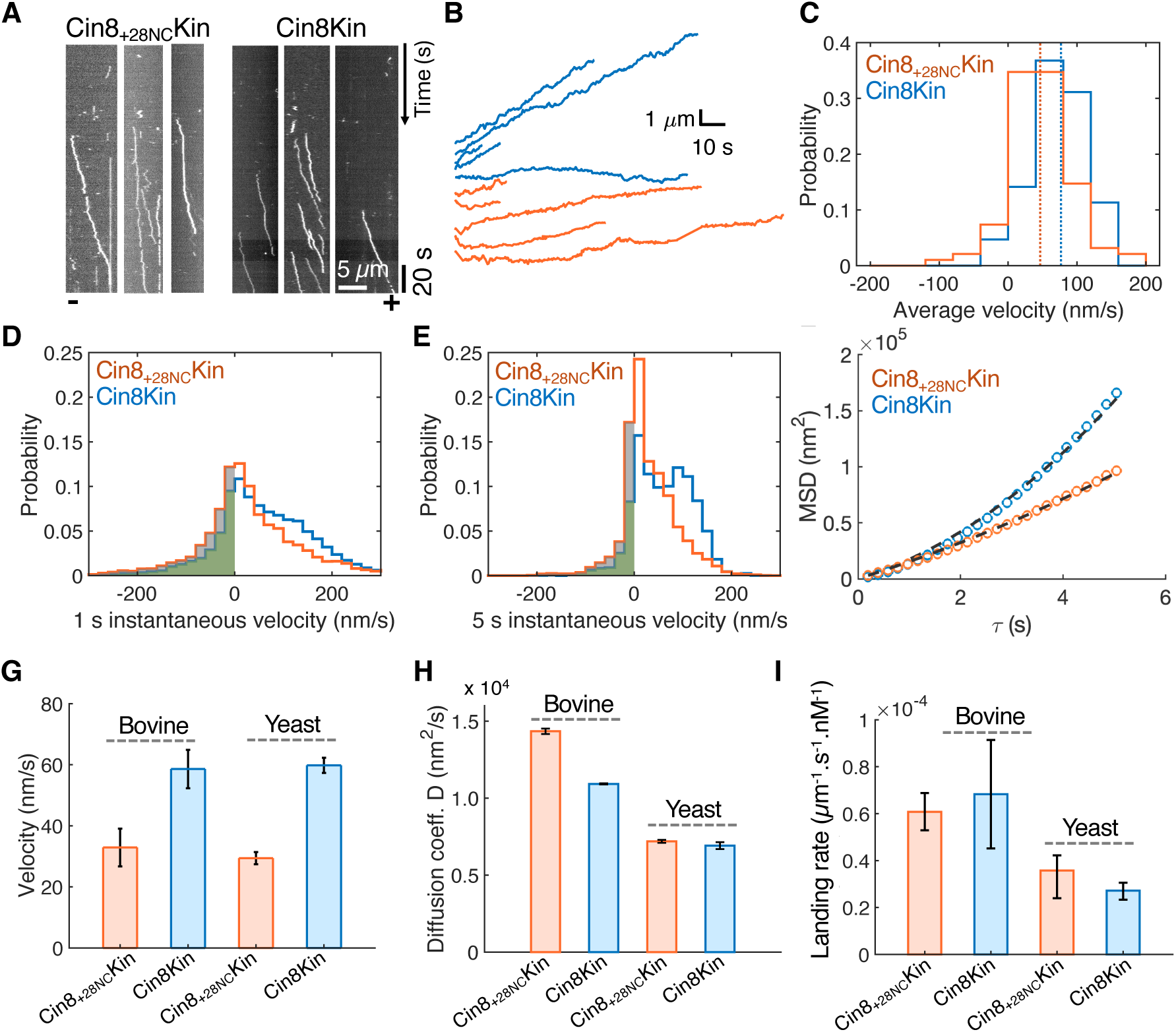
Cin8 dimer motility on yeast microtubules. **(A)** Representative kymographs and (**B)** displacement versus time traces for Cin8_+28NC_Kin (orange) and Cin8Kin (blue) motility in the presence of 1 mM ATP on yeast microtubules. Microtubule polarities are marked at the bottom of kymographs. **(C)** Distribution of whole trace velocities, with mean velocities 46.7 ± 4.4 for Cin8_+28NC_Kin and 76.6 ± 4.2 nm/s for Cin8Kin (mean ± SEM for 95 and 106 traces). **(D)** 1 s and **(E)** 5 s instantaneous velocity distributions. Gray (Cin8_+28NC_Kin) and green (Cin8Kin) shaded areas represent minus-end directed velocity segments. Average velocities from the 5 s velocity distributions were 29.4 ± 2.0 nm/s for Cin8_+28NC_Kin and 59.8 ± 2.5 nm/s for Cin8Kin (mean ± SEM for N = 726 and 179 independent 5 s windows from 95 and 106 independent molecules, respectively). **(F)** Mean squared displacement (MSD) analysis in ATP. Data were fit to, MSD = v^2^τ^2^+2Dτ (black lines), with velocities taken from Fig. 3E, yielding diffusion coefficients of 7194 ± 94 nm²/s and 6915 ± 220 nm²/s, respectively (fit ± 95% CI). Fit to MSD = aτ ^α^ + b yielded α of 1.29 for Cin8_+28NC_Kin and 1.64 for Cin8Kin (Table S1). (**G)** Comparison of velocities (summarized from Figs. 2E & 3E). (**H)** Comparison of diffusion coefficients and **(I)** motor landing rates for Cin8_+28NC_Kin (0.06 ± 0.007 and 0.036 ± 0.006 <u>μ</u>m^−1^ s^−1^ nM^−1^ for bovine vs yeast, respectively; mean ± SEM) and Cin8Kin (0.068 ± 0.02 and 0.027 ± 0.002 <u>μ</u>m^−1^ s^−1^ nM^−1^ for bovine vs yeast, respectively; mean ± SEM) on bovine and yeast microtubules (2 and 3 fields, respectively).

As an alternative way to quantify the degree of undirected motility, we carried out a mean squared displacement (MSD) analysis using the mean velocity from the instantaneous velocity distributions along with a diffusive term (Fig. 2F). Consistent with the wider instantaneous velocity distribution, the apparent 1D diffusion coefficient was 14338 ± 172 nm²/s for Cin8_+28NC_Kin and 10921 ± 40 nm²/s for Cin8Kin (fit ± 95% confidence interval, CI). Fitting to a model MSD = aτ^α^ + b, where α = 1 denotes pure diffusion and α = 2 denotes processive directional movement gave α of 1.29 for Cin8_+28NC_Kin and 1.38 for Cin8Kin (Table S1). Thus, consistent with its broader velocity distribution, Cin8_+28NC_Kin displayed more diffusion-like undirected movement than Cin8Kin.

Previous work on tetrameric Cin8 found that weakening motor-microtubule interactions by increasing the buffer ionic strength resulted in a shift to minus-end directionality (3,4,23). To test whether increasing ionic strength caused a similar directionality shift for Cin8 dimers, we repeated the experiments in BRB80 buffer supplemented with 110 mM KCl. In contrast to tetramers, clear net plus-end motility of Cin8_+28NC_Kin and Cin8Kin dimers was retained in the presence of added KCl, with slightly higher whole trace velocities of 47.3 ± 7.0 and 84.1 ± 7.6 nm/s, respectively (mean ± SEM) (Fig. S4). Thus, engineered Cin8 dimers display more inherent plus-end bias than native tetramers. No processive motility was observed in BRB80 buffer supplemented with 150 mM KCl (Fig. S4).

### Cin8 dimers move more directionally on yeast microtubules

To test the role of the microtubule track on Cin8 motility, we carried out single-molecule assays using recombinant *S. cerevisiae* tubulin, matching the species of the *S. cerevisiae* Cin8. To the best of our knowledge, only one previous study investigated the motility of Cin8 on yeast microtubule in vitro, and that study used full-length Cin8 tetramers from yeast lysates (43). On yeast microtubules, both Cin8 dimer constructs exhibited similar net plus-end directed motility as on bovine microtubules, though by visual inspection there were fewer fluctuations in directionality on the yeast microtubules (Fig. 3ACB; see also Fig. S5A). To quantify this undirected movement, we compared the fraction of traces that showed clear processive movements, defined as traces with average velocities >20 nm/s and net displacements >1 µm. Compared to bovine microtubules, the overall fraction of directed traces on yeast microtubules increased from 28% to 41% for Cin8_+28NC_Kin and from 26% to 68% for Cin8Kin (Fig. S5B). Similar to the bovine microtubules, replacing the proximal neck-coil of Cin8 with the stable kinesin-1 dimerization domain increased the mean trace velocity from 46.7 ± 4.4 nm/s for Cin8_+28NC_Kin to 76.6 ± 4.2 nm/s for Cin8Kin (mean ± SEM for 95 and 106 traces, respectively; Fig. 3C).

To quantify the relative proportions of directional and nondirectional movements on yeast microtubules, we carried out instantaneous velocity and mean squared displacement analyses. The instantaneous velocity analyses (Fig. 3DCE) revealed clear minus-end movements, particularly for Cin8_+28NC_Kin – from the 5 second instantaneous velocity distributions, 26.8% of segments were minus-end directed for Cin8_+28NC_Kin versus 13.7% for Cin8Kin (shaded regions). Cin8_+28NC_Kin also had a broader instantaneous velocity distribution than Cin8Kin with a standard deviation of 53.9 nm/s versus 33.4 nm/s, respectively (Fig. 3E). From the MSD analysis (Fig. 3FCH), the apparent diffusion constants on yeast microtubules were also similar to one another (7194 ± 94 nm²/s for Cin8_+28NC_Kin and 6915 ± 220 nm²/s for Cin8Kin); however fitting to MSD = aτ^α^ + b gave α of 1.29 for Cin8_+28NC_Kin and 1.64 for Cin8Kin, consistent with a greater degree of processive movement for Cin8Kin (Table S1). Interestingly, despite the shift toward more consistent plus-ended motility on yeast microtubules, motor landing rates on yeast microtubules were lower than on bovine microtubules (Fig. 3I). Overall, on yeast microtubules the mean velocities for each motor matched those on bovine microtubules (Fig. 3G); however, there was less undirected motility for both motors on yeast microtubules, which revealed more clearly the greater degree of undirected motility of Cin8_+28NC_Kin relative to Cin8Kin.

### Instantaneous velocity distributions are not consistent with simple diffusion

To gain further insight into the degree to which the undirected movements reflected bidirectional stepping versus thermally driven diffusion, we analyzed the instantaneous velocity distributions using a Gaussian Mixture Model. We first asked whether the velocity distributions on yeast microtubules could be decomposed into a sum of two normal distributions representing processive stepping and undirected diffusion or bidirectional stepping. A sum of two Gaussian did not fit the data well (Fig. S6). Instead, the velocity distributions for both Cin8Kin and Cin8_+28NC_Kin were well fit by the sum of three Gaussians (Fig. 4ACB). For each variant there was a principal mode (dominant peak) corresponding to plus-ended stepping; there was a narrow mode centered around zero (zero-centered peak), and there was a third mode with a very wide distribution (broad peak). The first difference between the two dimers was that the dominant velocity peak, which accounted for approximately 50% of the total distribution in both cases, exhibited markedly different mean velocities: 33.5 nm/s for Cin8_+28NC_Kin compared to 93.2 nm/s for Cin8Kin. The second difference was the width of the broad velocity peak, with Cin8_+28NC_Kin having a larger standard deviation (80.2 nm/s) than Cin8Kin (69.5 nm/s) and accounting for a greater proportion of the distribution (34% versus 24%). Together, these fits reiterate the observation that Cin8_+28NC_Kin exhibited greater stepping variability and reduced net plus-end velocity than Cin8Kin. In principle, the narrow modes centered around zero could reflect unbiased thermal diffusion while the broad peaks could represent bidirectional stepping with a plus-end bias. However, these data alone are not sufficient to support that interpretation. In sum, the presence of three modes in the velocity distribution argues against model consisting only of plus-end stepping and simple diffusion.

As a second test of whether simple diffusion can account for the observed undirected movements, we carried out stochastic simulations of Cin8 motility that included motor stepping and diffusion. We took the stepping rate from the mean velocity of the 5 s velocity distribution for each motor and added a diffusive term (〈x(t)〉^2^ = 2Dt), where D is the apparent diffusion constant. We generated simulated motility traces and analyzed them in the same manner as the experimental data to generate a 5 second instantaneous velocity distribution (Fig. 4 CCD). When we chose a value of D from the MSD analysis in ATP (7194 nm^2^/s and 6915 nm^2^/s), the resulting distributions were wider than the experimental and they also failed to capture the height of the peaks (cyan curves in Fig. 4 CCD). For both variants, a smaller D was able to reproduce the tails of the distribution, but the peaks were not aligned with the experimental peaks (purple curves in Fig. 4 CCD)). Finally, a larger D failed to capture either the peak or the tails of the distribution (magenta curves in Fig. 4 CCD). The failure of a simple plus-end directed stepping plus diffusion model to recapitulate the experimental data, together with the finding that the instantaneous velocity distributions were fit by three Gaussians (Fig. 4 ACB) argue against a model in which the undirected movements are solely due to thermally driven diffusion.

**Figure 4:**
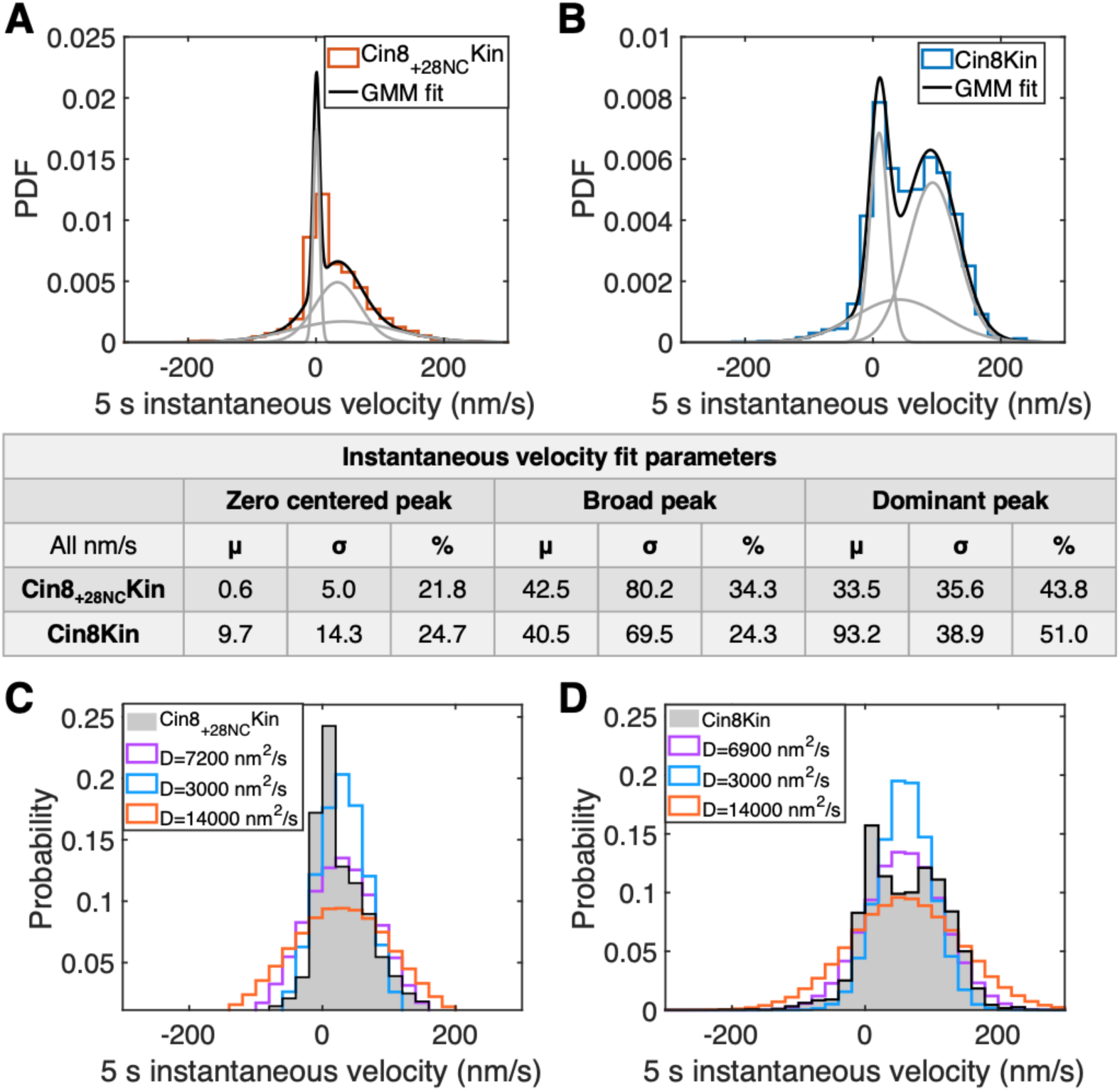
Separating diffusion from bidirectional stepping. **(A and B)** Gaussian Mixture Model analysis of 5 s instantaneous velocity distributions. Velocity distributions on yeast microtubules (from Fig. 3E) were fit by three Gaussians (gray lines), with values given in table. (**C and D**) Comparison of experimental results with simulations of a motor with a unidirectional velocity plus random diffusion. Motor velocities were taken from mean of instantaneous velocity distributions in Fig. 3E. Initial diffusion constants (purple line) were taken from MSD analysis in Fig. 3F and varied up and down by two-fold. The experimental velocity distribution from Fig. 3DCE are shown in gray, whereas the distributions shown by cyan, purple and magenta represent simulated data with diffusion coefficients values given in figure inset. In the table, µ, σ and % represent mean, standard deviation, and weight of each Gaussian distribution, respectively.

### ATPase kinetics indicate that flexibility of the Cin8 neck coil diminishes mechanochemical coupling

Understanding how ATP hydrolysis is coupled to mechanical stepping is central to understanding the kinesin stepping mechanism. For kinesin-1, the stepping rate (k_step_) matches the ATP hydrolysis rate (k_cat_), consistent with the motor hydrolyzing one ATP per step, defined as tight mechanochemical coupling (44,45). In our single-molecule assays, both Cin8 dimers showed net plus-end directed processive motility along with undirected fluctuations, which in principle could result from either pure diffusion with no ATP hydrolysis (the consensus model for processive monomeric KIF1A motors (46)) or from bidirectional stepping (with each step consuming one or more ATP). Cin8_+28NC_Kin, which contains its native proximal neck-coil domain, had a slower net plus-end velocity and more undirected movement. If this undirected movement was simply diffusion, the Cin8_+28NC_Kin ATPase rate would be expected to be lower than that of Cin8Kin, reflecting the time the motor spends in a diffusive state. In contrast, if the greater undirected movement of Cin8_+28NC_Kin was due to bidirectional stepping, the ATPase rate would be expected to be higher than Cin8Kin. To examine the degree of mechanochemical coupling for the two Cin8 dimers, we measured the microtubule-stimulated ATPase rates at varying concentrations of bovine microtubules using an NADH-coupled assay (Fig. 5A). Cin8Kin had a mean plus-end velocity of 58.6 nm/s on bovine microtubules (Fig. 2E), corresponding to 7 steps/s, and the measured k_cat_ was 5.1 ± 1.5 s⁻¹ (± 95% CI; Fig. 5B). This agreement is consistent with tight coupling between ATP hydrolysis and stepping. However, the data are also consistent with the motor sharing time between a diffusive state with no ATPase and bidirectional stepping state with an elevated ATPase rate. In contrast, Cin8_+28NC_Kin had a slower mean plus-end velocity of 23.9 ± 6.9 nm/s on bovine microtubules (Fig. 2E), corresponding to 3 steps/s, and a higher k_cat_ of 6.9 ± 2.3 s⁻¹ (± 95% CI). The ∼two-fold faster ATPase rate for Cin8_+28NC_Kin compared to its stepping rate suggests that instability of the native Cin8 neck coil leads to uncoupling between stepping and hydrolysis. This uncoupling also suggests that the greater proportion of undirected movement for Cin8_+28NC_Kin (Fig. 2 & 3) results from bidirectional stepping and not from enhanced diffusion.

**Figure 5:**
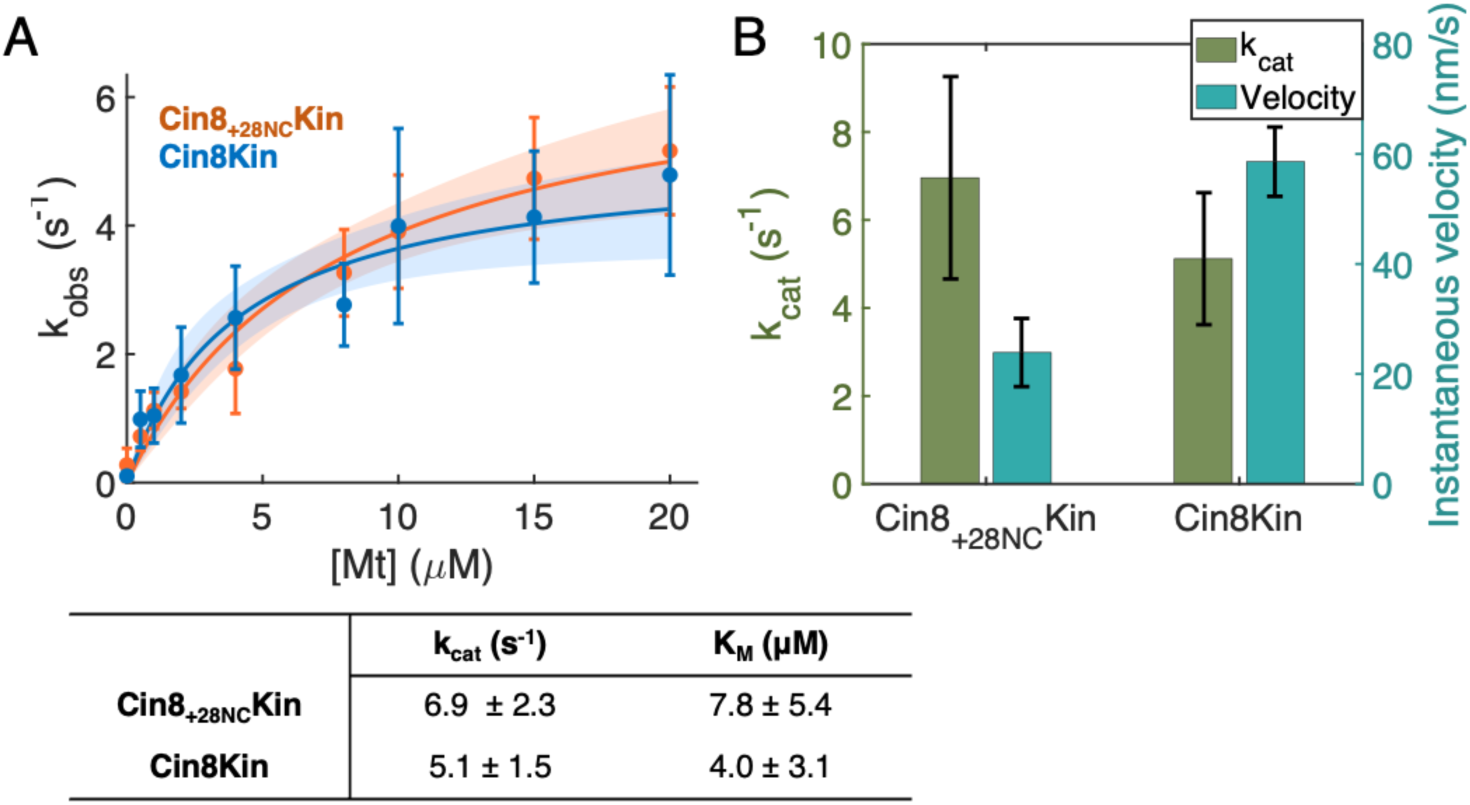
ATP turnover kinetics and corresponding stepping velocities. **(A)** Microtubule-stimulated ATP hydrolysis of Cin8 dimers in BRB80 buffer at various bovine microtubule concentrations (n = 5 trials per point). Data were fit to the Michaelis-Menten equation, using 1/SEM weighing for each point, to obtain the k_cat_ and a K_m_ (± 95% CI) presented in the table. Shaded area represents fit ± 95% CI. **(B)** Comparison of k_cat_ (fit ± 95% CI; left axis, green) to the 5 s mean instantaneous velocity (mean ± SEM; right axis, blue) for Cin8_+28NC_Kin and Cin8Kin (from Fig. 2D).

### Cin8 dimers diffuse on MTs for long durations in the ADP-bound state

In a further effort to separate out thermally-driven diffusion from bidirectional stepping, we examined the behavior of the Cin8 dimers on yeast microtubules in the presence of 1 mM ADP, which generates a diffusive weak-binding state. Unlike most kinesins, which bind weakly to microtubules in ADP (28,47,48), Cin8 tetramers have been shown to bind for long durations in ADP (3,23,25,26). We found that both engineered Cin8 dimers remained bound to microtubules for long durations in ADP (Fig. 6A). Thus, consistent with previous observations (24), Cin8 motor domains have inherently high microtubule affinity in ADP independent of their tetrameric structure or their tail domains.

**Figure 6:**
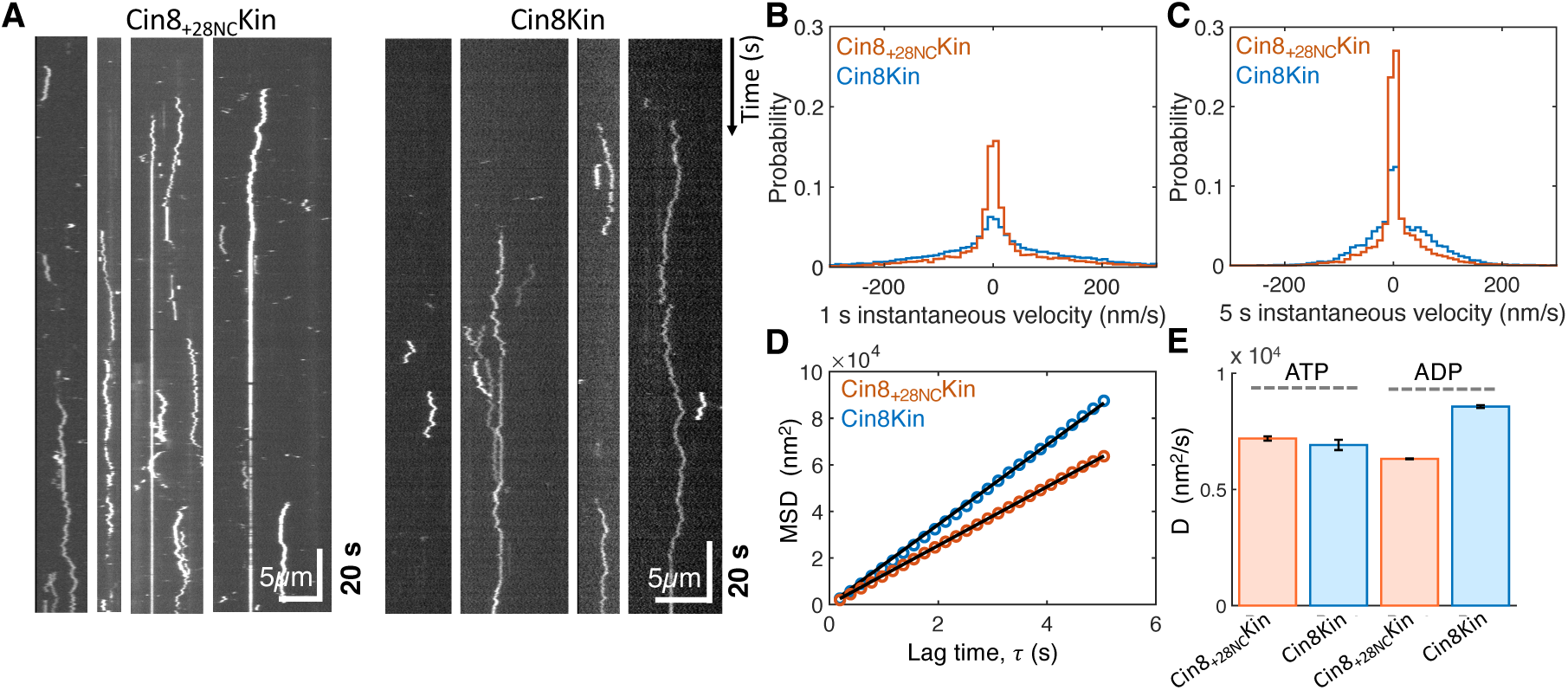
Cin8 remains bound to microtubules in the presence of ADP. **(A)** Representative kymographs in the presence of 1 mM ADP on yeast microtubules. **(B)** 1 s and **(C)** 5 s instantaneous velocity distributions for Cin8_+28NC_Kin (orange) and Cin8Kin (blue) (from N = 59 and 97 traces, respectively). **(D)** Mean squared displacement (MSD) comparison in ADP. Data were fit to MSD = 2Dτ, yielding D = 6314 ± 17 nm^2^/s for Cin8_+28NC_Kin and 8565 ± 58 nm^2^/s for Cin8Kin (fit ± 95% CI). **(E)** Comparison of apparent diffusion coefficients for Cin8_+28NC_Kin and Cin8Kin in ATP and ADP on yeast microtubules.

To quantify motor diffusion in the ADP-bound state, we plotted the instantaneous velocity distributions for both motors and carried out a mean squared displacement (MSD) analysis. Both 1 s and 5 s instantaneous velocity distributions are centered around zero, as expected, indicating that directional motility is ATP hydrolysis-dependent (Fig. 6BCC). The apparent diffusion constants in ADP were 6314 ± 17 nm^2^/s for Cin8_+28NC_Kin and 8565 nm^2^/s ± 58 nm^2^/s Cin8Kin (fit ± 95% CI; Fig. 6DCE), which are in the same range of the corresponding diffusion constants in ATP on yeast microtubules (Fig. 6E). Thus, the data don’t exclude the possibility that the nondirected movements in ATP are 1D diffusion of the motors during phases when the motors are not stepping.

### Cin8 neck-coil domain enhances microtubule binding duration in ATP but not ADP

To investigate the influence of inter-head tension on the overall microtubule affinity of Cin8_+28NC_Kin and Cin8Kin, we quantified the residence times of Cin8 dimers in ATP and ADP on yeast microtubules. If flexibility of the Cin8 neck-coil reduces inter-head coordination needed for processive stepping, then Cin8_+28NC_Kin should have shorter binding durations than Cin8Kin. In contrast, if flexibility of the Cin8 neck-coil reduces steric constraints and allows both heads to bind to the microtubule more readily, then Cin8_+28NC_Kin should have longer binding durations than Cin8Kin. Mean binding durations in ATP were calculated by fitting an exponential function to the distribution of run durations. We found that Cin8_+28NC_Kin remained bound to microtubules ∼3-fold longer than Cin8Kin (26.3 ± 0.32 s and 8.9 ± 0.27 s, respectively (fit ± 95% CI); Fig. 7A). Thus, the data are consistent with flexibility of the Cin8 neck coil reducing steric constraints between the two heads and promoting a two-head binding state in ATP. To test whether similar behavior occurs in ADP, we repeated the analysis in 1 mM ADP and found the opposite result, with the Cin8_+28NC_Kin binding duration (10.6 ± 0.61 s) being shorter than that for Cin8Kin (15.8 ± 0.67 s; Fig. 7B & C). Thus, in ATP the proximal Cin8 neck domain prolongs the microtubule binding duration in ATP, whereas in ADP it shortens the binding duration.

**Figure 7:**
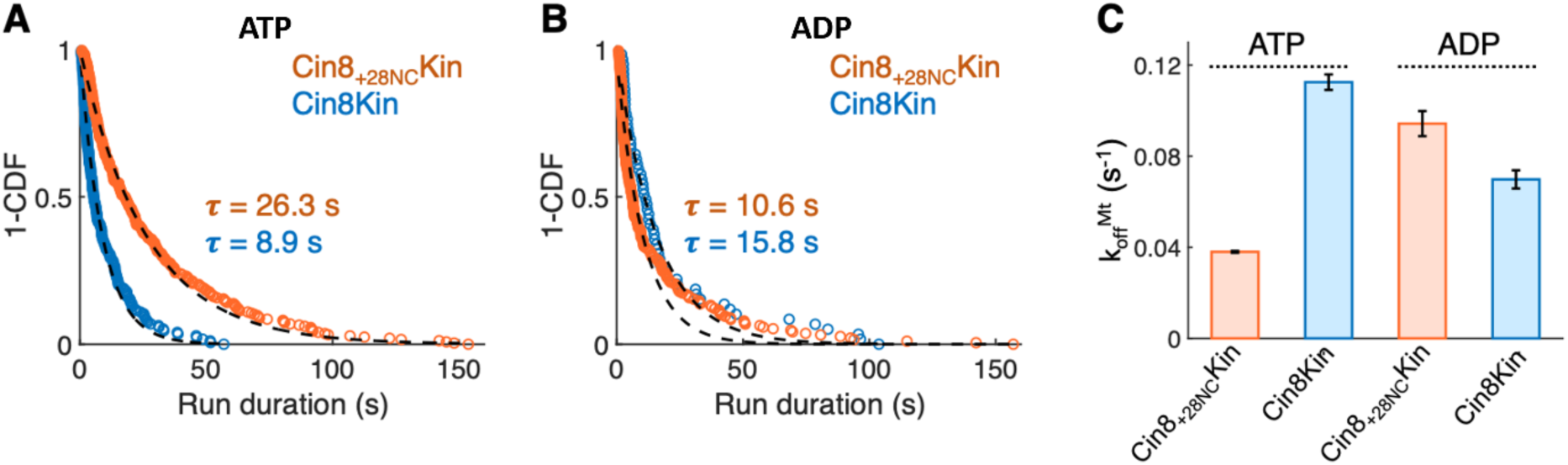
Microtubule binding durations of Cin8 dimers on yeast microtubules. Distribution of bound durations (τ) from single-molecule motility traces of Cin8_+28NC_Kin and Cin8Kin in **(A)** ATP and **(B)** ADP. Data are plotted as 1 - Cumulative distribution function (1-CDF) and fit with an exponential (black lines). **(C)** Comparison of microtubule detachment rates (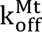), calculated as 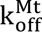 = 1/τ. Sample sizes (n) for Cin8_+28NC_Kin were 246 (ATP) and 190 (ADP); for Cin8Kin, 172 (ATP) and 59 (ADP). Error bars represent 95% CI. Processive and diffusive traces in ATP were analyzed separately and found to have similar binding durations (Fig. S7).

## Discussion

For transport kinesins, processivity relies on mechanochemical coordination between the two motor domains; however, the degree to which the two heads of Cin8 are coordinated and whether this coordination plays a role in the bidirectional motility of Cin8 are not clear. We find that engineered Cin8 dimers move with a plus-end bias and exhibit undirected movements that result from a combination of ATP-driven bidirectional stepping and thermally-driven diffusion. Including the native neck-coil of Cin8 leads to more undirected movements and a slower net plus-end speed, no change in the ATP turnover rate, and a longer residence time on the microtubule. On yeast microtubules, Cin8 moved at similar speeds as on bovine microtubules but with fewer undirected movements. These results suggest that, in contrast to kinesin-1, plus-ended movement of Cin8 requires little coordination between the activities of its motor domains.

### Cin8 dimer motility is consistent with previous work

Motility of the two Cin8 dimer variants studied here is broadly consistent with previous work. Duselder et al constructed a Cin8Kin containing the full N-terminal, 4 residues more in the neck domain, and a shorter kinesin-1 dimerization domain (24). On porcine microtubules in BRB80 buffer pH 6.8 without added salt, the dimer showed bidirectional movements, a small plus-ended mean velocity (18 nm/s), and an MSD exponent of 1.24; behavior that is consistent with directional motility. Our Cin8Kin dimer had a faster net velocity (62 nm/s on bovine microtubules from Fig. 2E), a slightly higher MSD exponent of 1.38, and clear plus-end directed segments not observed in that work. Although there are differences in the motility analysis between the two studies, differences in the truncation points of the constructs are the most likely explanation for the differences in motility. Consistent with this, previous work using dimers having the same motor domain length as our Cin8Kin (38–530) with a different dimerization strategy found similar plus-end gliding velocities in multi-motor gliding assays (25). One consistent result from these dimer studies was that the fast minus-end motility seen in Cin8 tetramers was absent, consistent with regions outside the motor domain being required for fast minus-end velocities seen in tetramers.

### Diffusion and bidirectional stepping are not easily separated

Because bidirectional stepping is a form of a random walk and random walks inherently generate diffusion-like behavior (49), it is challenging to separate out contributions of ATP-driven bidirectional stepping from simple thermally-driven diffusion. The first argument in favor of bidirectional stepping is the substantial fraction of minus-end velocity segments in the instantaneous velocity distributions (Fig. 2E and 3E). When we analyzed whether a model consisting of plus-ended stepping together with thermal diffusion could recapitulate the instantaneous velocity histograms, we failed to find a diffusion constant that matched both the height of the velocity peak and the width of the overall distribution. Although we can’t rule out the possibility that more complex stepping and diffusion models could better match the data, our analysis argues against diffusion alone accounting for the undirected motility.

The second argument supporting bidirectional stepping is the fit of a three-component Gaussian Mixture Model to the instantaneous velocity histograms (Fig. 4A). In all cases the fits included a dominant peak (roughly half of the total) centered around the mean plus-end velocity, consistent with active stepping. There was also a peak near zero (∼20-25% of total), consistent with thermally-driven diffusion. The third peak was broad and centered at a positive velocity, properties consistent with it representing bidirectional stepping with a plus-end bias. The fact that this broad peak represented 34% of the data for Cin8_+28NC_Kin but only 24% for Cin8Kin is consistent with the native Cin8 neck coil leading to more bidirectional stepping.

The third argument supporting bidirectional stepping as the dominant source of undirected movement is the relationship of the motor velocities to their ATPase rates. For Cin8Kin, the ATPase rate (k_cat_ of 5 s^−1^) was similar to the net stepping rate calculated from the mean velocity (7 s^−1^). This match could be interpreted as ATP hydrolysis being tightly coupled to forward stepping; however, the substantial undirected movements suggest a more complicated picture involving some combination of bidirectional stepping and diffusion. The more important result was that the k_cat_ of Cin8_+28NC_Kin (7 s^−1^) was more than twice the net stepping rate (3 s^−1^). This discrepancy suggests that the observed net velocity results from a combination ATP-driven plus- and minus-end directed steps. For instance, if one ATP was associated with every step, the results could be explained by a model with a 5 s^−1^ plus-end stepping rate and a 2 s^−1^ minus-end stepping rate. However, the undirected movements we observed, which lasted for multiple seconds for some motors and the entire duration for a subset of motors, suggest that the Cin8 dimers alternate between phases of ATP-driven stepping and thermally-driven diffusion where little or no stepping occurs. In this case, diffusive periods with zero ATPase could be balanced by periods of bidirectional stepping at rates substantially above the average. Inferring these details from ensemble averaged ATPase rates is speculative; the notable result is that Cin8_+28NC_Kin had a higher ATPase rate and a lower net stepping rate than Cin8Kin.

Two pieces of evidence argue that at least some portion of the undirected movements reflect thermally-driven diffusion. The first is that when comparing the behavior in ATP versus ADP, both the apparent diffusion constants and the microtubule off-rates differ very little (Fig. 6E and 7C). Because the ADP state is generally the lowest affinity state in the kinesin ATP hydrolysis cycle (47,50), both the diffusion constant and the dissociation rate would be expected to be highest in ADP. One interpretation is that Cin8 spends most of its ATP hydrolysis cycle in the ADP state, similar to what has been suggested for the superprocessive kinesin-3 motor, KIF1A (28,51). Unfortunately, the high affinity of Cin8 for mant-ADP (Fig. S8) prevented us from measuring the ADP off-rate directly. The second argument that diffusion makes up part of the undirected movement is the different behaviors on yeast versus bovine microtubules. Consistent with the general conservation of tubulin from fungi to mammals, the mean velocities observed on the two substrates were similar for both motors. However, there were more undirected movements on bovine microtubules than on yeast microtubules, particularly for Cin8Kin. Motor landing rates were also higher on bovine microtubules. We tentatively interpret the enhanced undirected movements on bovine microtubules as resulting from greater electrostatic interactions between Cin8 and the bovine C-terminal tails, which shift the motor from a stepping state to a diffusive state. The importance of the C-terminal tail to Cin8 motility was demonstrated by a recent study that found that deleting the yeast β-tubulin C-terminal tail reduces the on-rate, run length and velocity of full-length Cin8 (43).

### Cin8 lacks tight inter-head coordination

In contrast to kinesin-1 where processivity is achieved by tight coordination between the mechanochemical cycles of the two heads, we favor a model for Cin8 in which there is little inter-head coordination and processivity is instead achieved by the inherent high affinity of the motor domains for the microtubule. We found that in the presence of ADP, Cin8 dimers bound to microtubules for long durations, consistent with previous observations for tetrameric, dimeric, and monomeric Cin8 (3–5,24,25). This property differs from the relatively fast microtubule off-rate of kinesin-1 in ADP state (47), and it relieves the constraint inherent in kinesin-1 that the mechanochemical cycles of the heads remain out of phase to minimize time in the ADP state (18). Notably, we found that Cin8_+28NC_Kin moved at a slower mean velocity than Cin8Kin and had more undirected motility, but it remained bound to the microtubule nearly three-fold longer than Cin8Kin (Fig. 7A). We interpret this to mean that the relative instability of the native Cin8 neck-coil domain diminishes mechanical coupling between the two motor domains, resulting in Cin8_+28NC_Kin spending a greater fraction of time in the two-heads-bound state than Cin8Kin. It was shown previously that disrupting the mechanical connection between the two heads of kinesin-1 by inserting six or more residues into the neck linker domain resulted in significant backstepping under hindering loads, longer residence times under no load, and the motor losing the requirement of ATP binding to occupy the two-heads-bound state (21,22). Thus, there is precedence for a loosening of the mechanical connection between the two heads of kinesin enabling the two-head-bound state.

Further evidence supporting a lack of inter-head coupling in Cin8 come from comparisons of our dimers to previous work on Cin8 monomers. It was found previously that in multimotor gliding assays, monomeric Cin8 motor domains (residues 1-528 linked to gelsolin for surface immobilization) moved microtubules at 20-40 nm/s, similar to our gliding speeds for dimeric Cin8_+28NC_Kin (Fig. 1) (52). Furthermore, in ATPase assays Cin8 monomers (residues 38-530) were found to have a k_cat_ of 4.1 s^−1^ (25), which is roughly half of our dimer ATPase rates (meaning the rates are the same on a per-head basis). Thus, multimotor gliding speeds and ATPase rates appear to be equivalent whether the heads are connected through the Cin8 neck-coil or not. And in dimeric and monomeric Cin8, plus-end directed and undirected motility appear to be the default modes. Notably, in Cin8Kin where we connected two Cin8 heads directly through the stable kinesin-1 neck-coil domain, the motor moved faster and with more plus-end directionality than when the Cin8 neck-coil was present. These results suggest that features that regulate Cin8 directionality, such as the tail domain (24), N-terminal extension (26), loop-8 (25) and phosphorylation (53,54) could exert their effects by altering inter-head coordination. Similarly, geometries that promote plus-end directionality such as motor clustering (4,23) and crosslinking of antiparallel microtubules (3,13) could be thought of as mechanically coordinating the motor domains.

## Author contributions

H.P., W.O.H. and L.G. developed original ideas for the project. H.P. and W.O.H. designed research. H.P. and T-C.M. carried out experiments and analyzed data. T-C.M. carried out simulations. H.P., T-C.M., L.G. and W.O.H. interpreted data. E.B. and L.M.R. generated yeast tubulin and helped design experiments. H.P. and W.O.H. wrote the manuscript and all authors edited the manuscript.

## Acknowledgments

This work was funded by NIH GM GM139568 awarded to W.O.H., NIH R35 GM156385 awarded to L.M.R., and ISF grant 629/23 awarded to L.G. The authors thank members of the Hancock and Gheber labs for helpful suggestions.

## Supporting information

**Figure S1:**
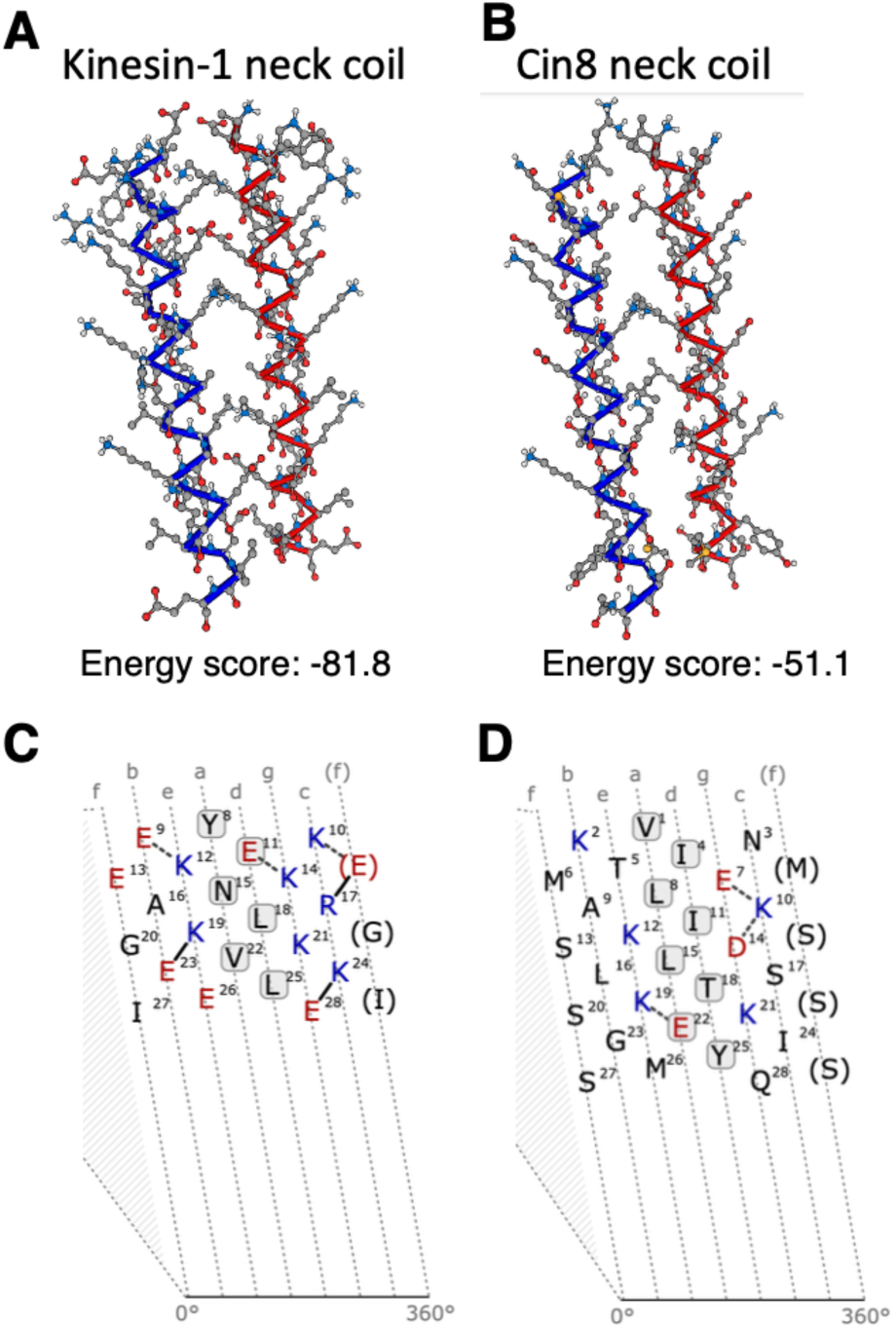
Cin8 is predicted to have a less stable neck coiled-coil than kinesin-1. **(A, B)** Coiled-coil prediction models for the proximal neck coil domains of kinesin-1 (A) and Cin8 (B) generated using CCbuilder 2.0 (1). The predicted coiled-coil stability scores were −81.8 for kinesin-1 and −51.1 for Cin8, with more negative values indicating greater stability. The backbones of the interacting helices are shown in blue and red, with side chains depicted in grey. **(C, D)** Predicted stabilizing interactions within the coiled-coil domains of kinesin-1 (C) and Cin8 (D). For kinesin-1, three strong (solid lines) and three intermediate (dashed lines) stabilizing interactions were identified, whereas only three intermediate interactions were predicted for Cin8. Coiled-coil predictions and alignment were performed using CCbuilder 2.0, COcoNat (2), T-Coffee (3), and Waggawagga (4).

**Figure S2:**
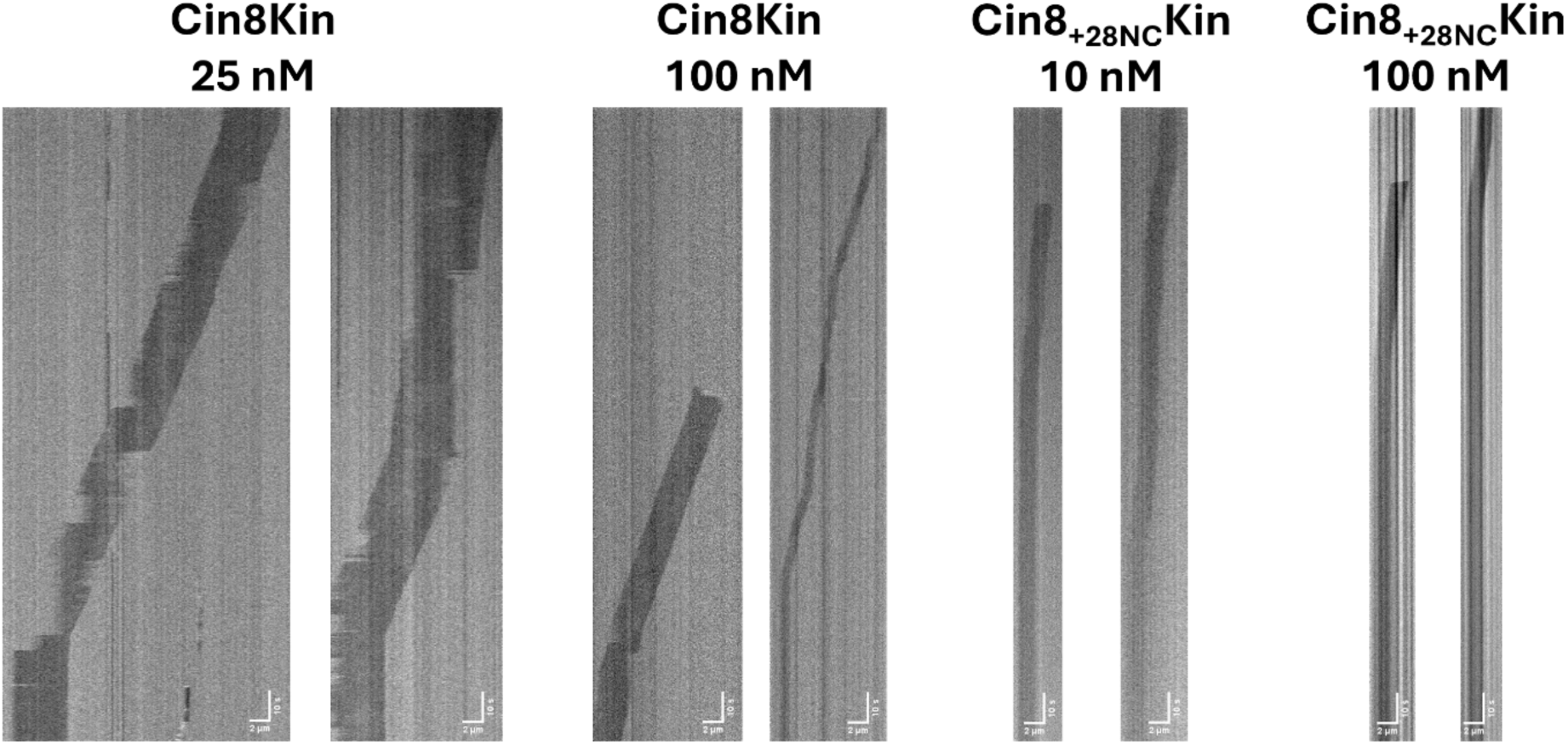
Kymograph of Cin8 dimer gliding assay. Example traces are shown at highest and lowest motor densities where motility was observed for each motor. Plus-ended motility was confirmed for all examples by washing in KIF1A-GFP at end of experiment.

**Figure S3:**
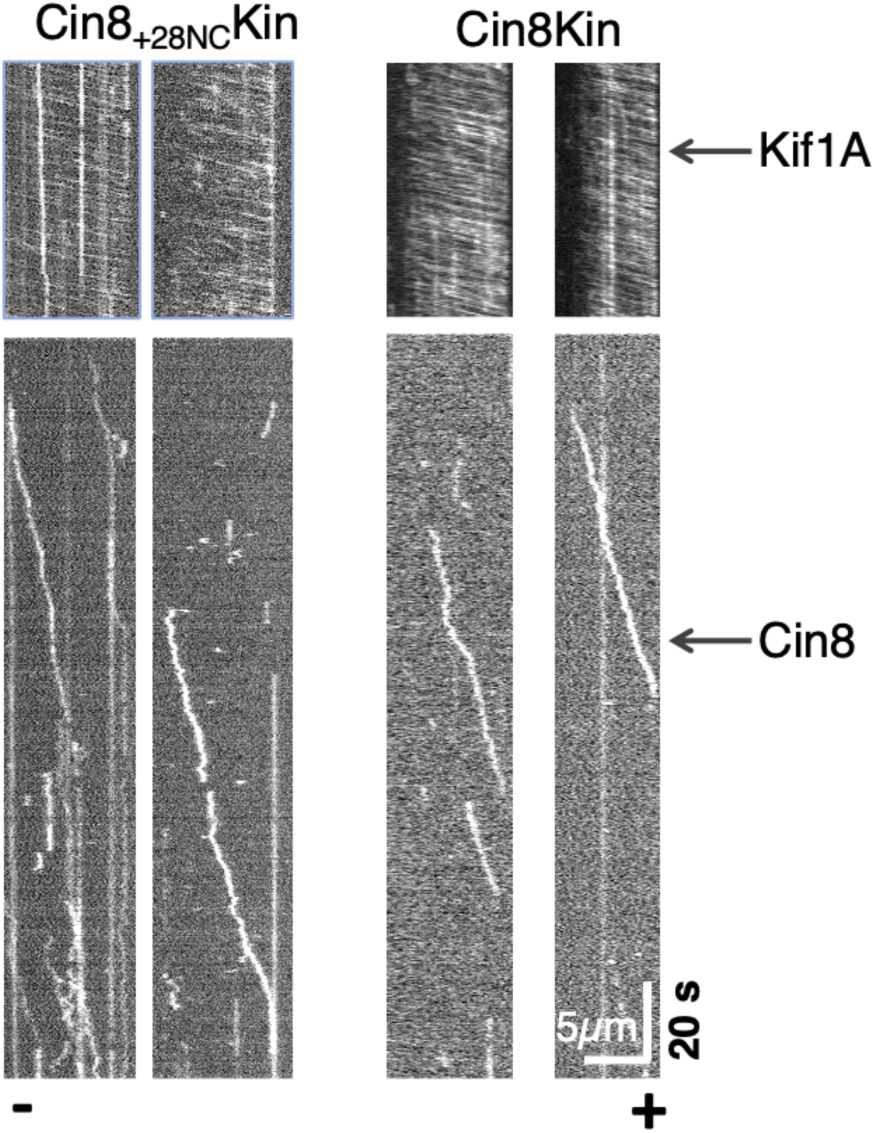
Microtubule polarity determination. To define microtubule polarity, fluorescent KIF1A (5) was introduced at the end of the experiment (top). Kymographs at bottom are same microtubule showing Qdot-labeled Cin8_+28NC_Kin or Cin8Kin motility, with microtubule polarity marked at the bottom of kymograph.

**Figure S4:**
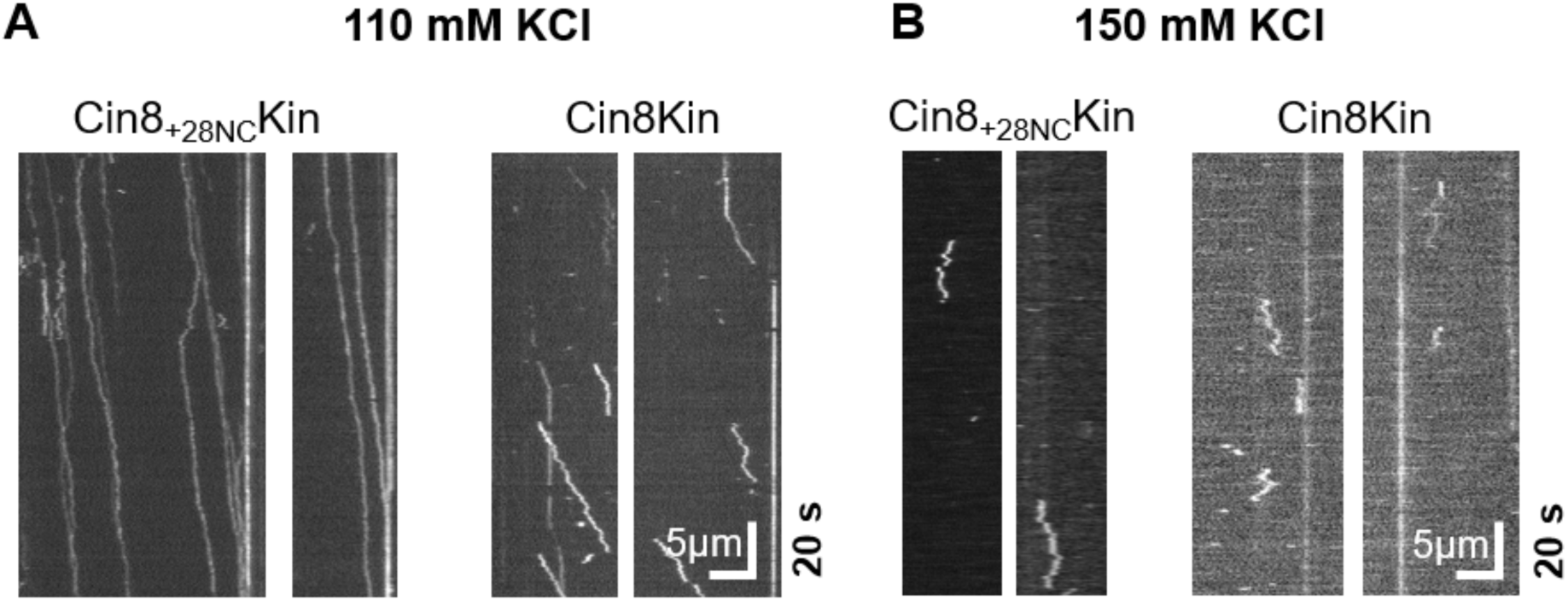
Single-molecule motility of Cin8 dimers on bovine microtubules in BRB80 buffer plus added salt. **(A)** Motility in BRB80 buffer adjusted to pH 7.2 plus 110 mM KCl. Net plus-end motility was retained even at elevated ionic strength with whole trace velocities of 47.3 ± 7.0 and 84.1 ± 7.6 nm/s for Cin8_+28NC_Kin (n = 27) and Cin8Kin (n = 30), respectively (mean ± SEM). **(B)** Motility in BRB80 buffer adjusted to pH 7.2 plus 150 mM KCl. Landing event frequency decreased drastically above 110 mM KCl and motors that landed have very short attachment times and no clear processive motility.

**Figure S5:**
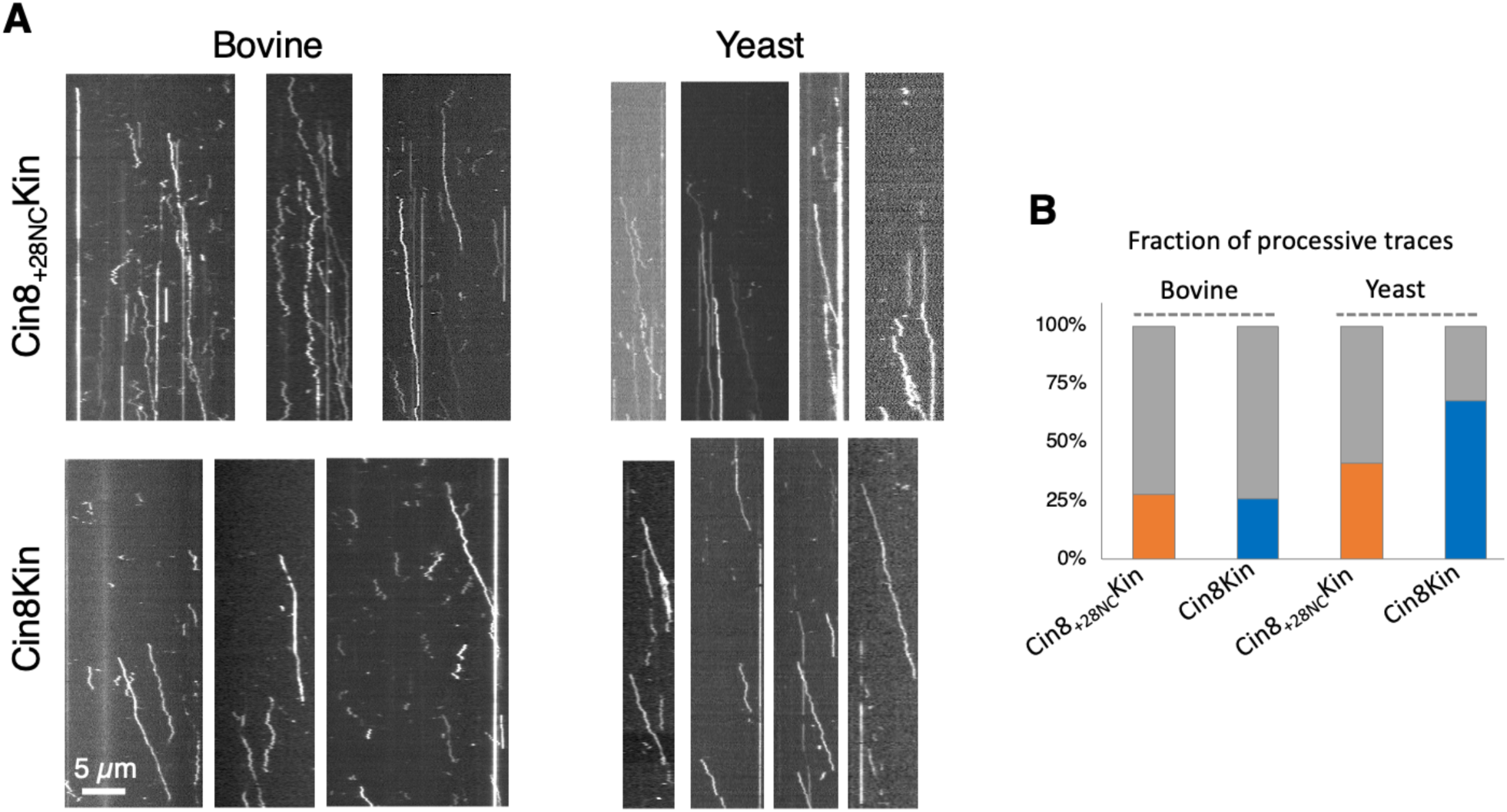
Comparison of Cin8 dimer motility on bovine and yeast microtubules. **(A)** Representative kymographs for Cin8_+28NC_Kin and Cin8Kin motility on bovine or yeast microtubules as indicated at the top. On yeast microtubules Cin8 dimers move with more consistent plus-end directionality. **(B)** Fraction of processive traces for each condition, defined as traces with overall velocity >20 nm/s and overall displacement >1 µm. The grey area at the top of each bar represents fraction of motors with undirected motility.

**Figure S6:**
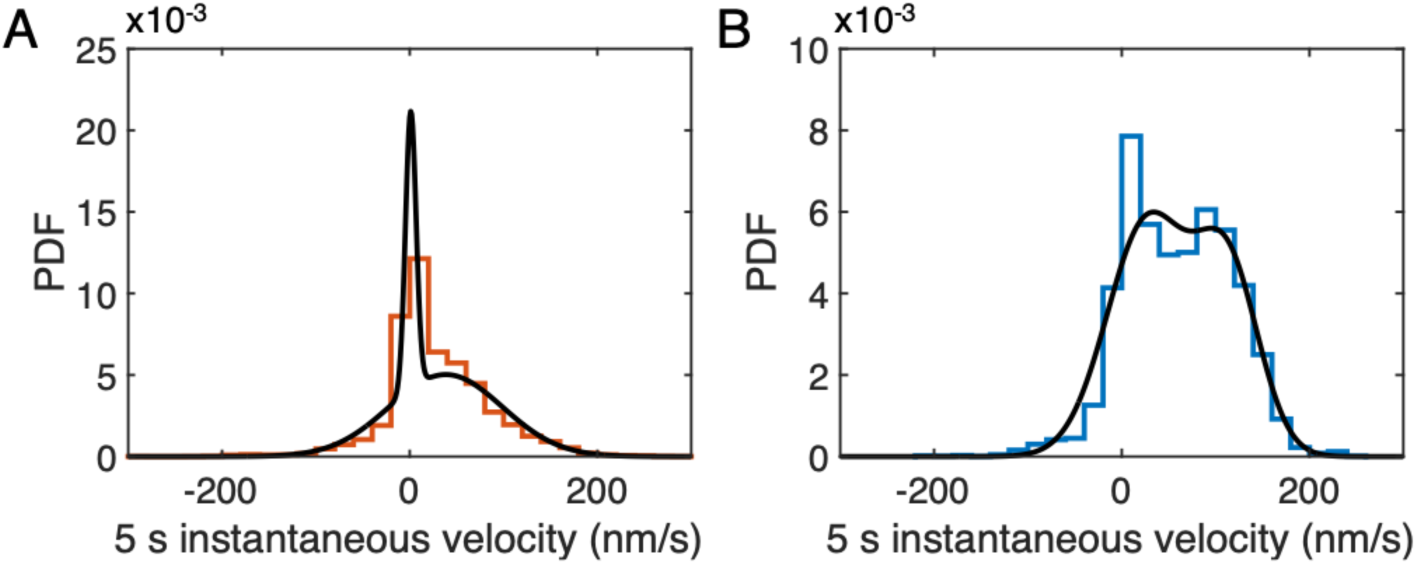
Velocity distributions for Cin8_+28NC_Kin (A) and Cin8Kin (B) could not be fit well by the sum of two Gaussians. The velocity distributions are depicted in orange (Cin8_+28NC_Kin) or blue (Cin8Kin) and the Gaussian Mixture Model (GMM) fit in black. The Bayesian Information Criterion (BIC) of 3-Gaussian and 2-Gaussian models for Cin8_+28NC_Kin are 1.01×10^5^ and 1.02×10^5^, respectively. For Cin8Kin, the BIC of 3-Gaussian and 2-Gaussian models are 9.29×10^4^ and 9.36×10^4^, respectively. These results indicate that the distributions are better explained by three Gaussians.

**Figure S7:**
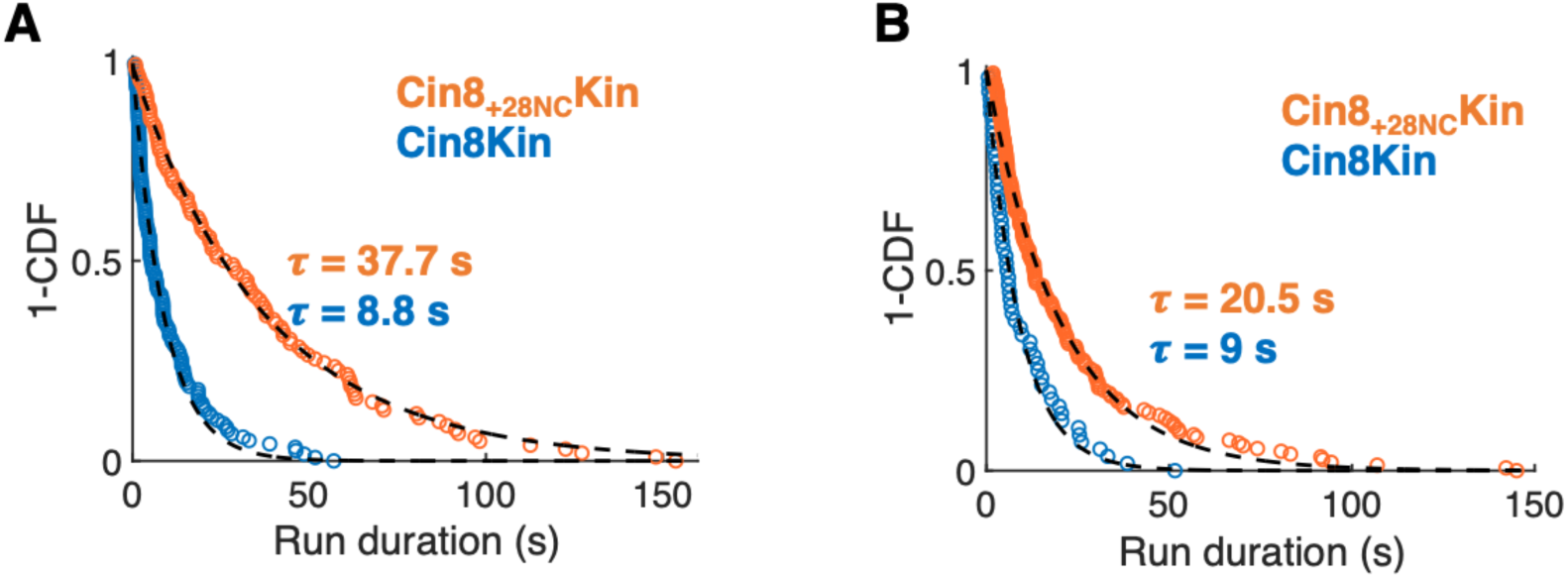
Microtubule binding durations of processive and undirected Cin8 dimers on yeast microtubules in ATP. Processive molecules were defined as those with average velocities >20 nm/s and net displacements >1 µm. Cumulative distribution functions (1-CDF) of run durations for **(A)** processive and **(B)** undirected motility traces for Cin8_+28NC_Kin and Cin8Kin were fit by single exponential functions to obtain the mean run duration, τ, for each.

**Figure S8:**
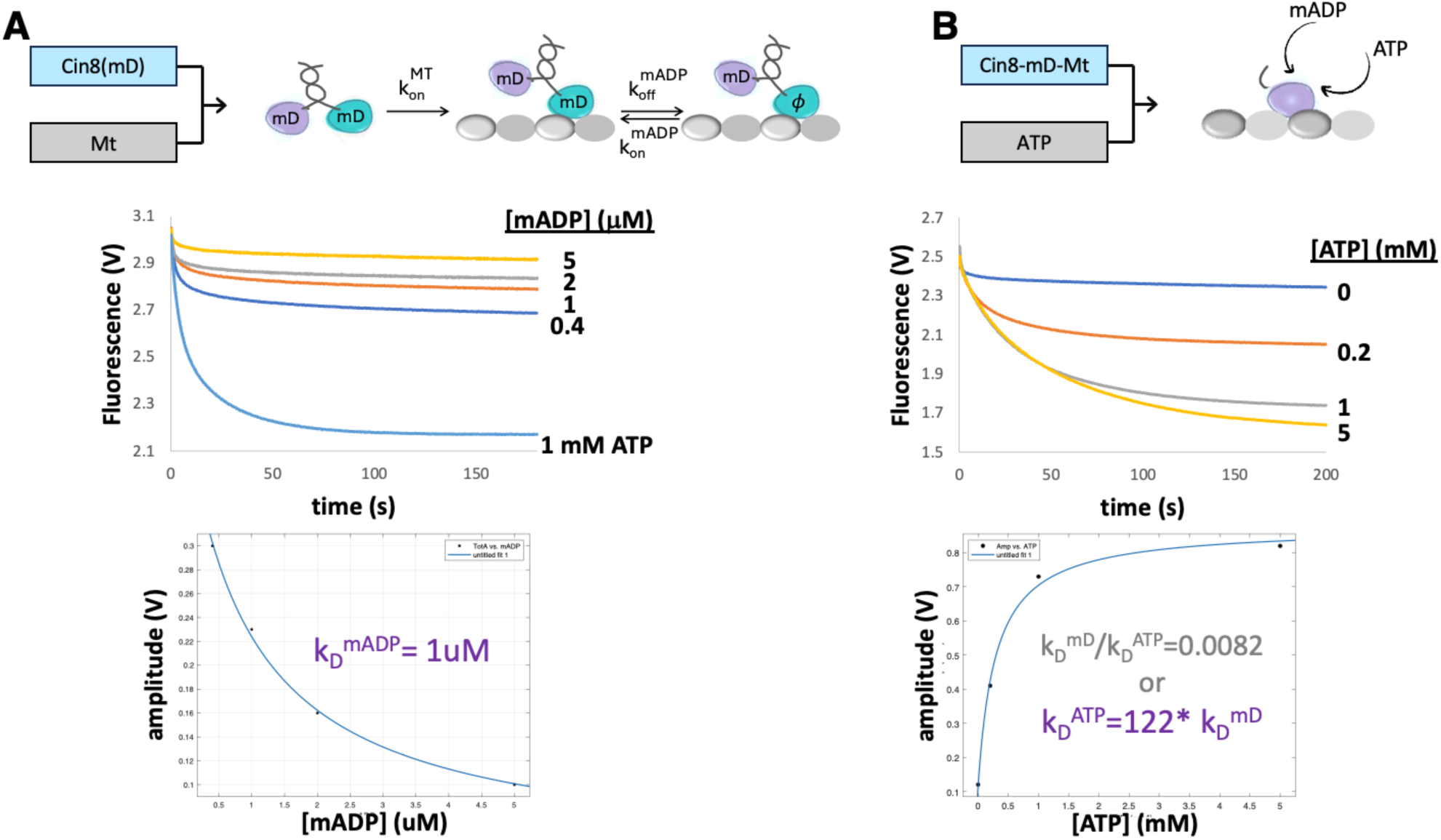
Cin8 has a high affinity for mant-ADP. **(A)** To measure microtubule-stimulated mant-ADP release, mant-ADP labeled Cin8Kin dimers were flushed against microtubules in the presence of varying concentrations of free mant-ADP. Experiments were carried out at 21 °C in BRB80 buffer using an Applied Photophysics SX20 spectrofluorometer at 356-nm excitation with a 395 long-pass emission filter. The conditions in the chamber are 5 µM microtubules, 0.2 µM Cin8, and mADP or ATP as noted. Note that the fall in fluorescence, corresponding to mantADP release is small and concentration dependent, denoting incomplete mantADP release. Full mantADP release was achieved by flushing mADP labelled motors against 5 µM microtubules + 1 mM ATP (chamber concentrations). To determine the mantADP affinity of microtubule-bound Cin8, the fall in the fluorescence amplitude was plotted against the free mant-ADP concentration and fit using 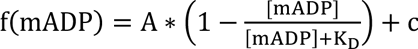, where A is the maximum amplitude, c is an offset, and K_D_ is the dissociation constant. The fit estimated 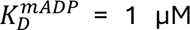 for microtubule-bound Cin8. **(B)** A competition assay between mant-ADP and ATP in the absence of microtubules was performed by flushing microtubule-bound mant-ADP labeled Cin8 monomer against varying concentrations of ATP. The amplitude of the fall in fluorescence is plotted against [ATP] and fit using f(ATP) = A ∗ 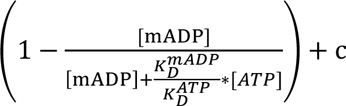, where A is the maximum amplitude and c is the offset. The resulting fit gave 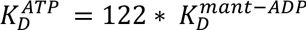. Thus, the affinity for mantADP is 122 times tighter than the affinity for ATP. This complicates mantADP stopped flow experiments with Cin8.

**Table S1.** Fits to MSD results in Figure 2F and 3F where α=1 represents diffusion and α=2 represents processive movement.

| Fit:<br>MSD = $a\tau^\alpha + b$ | Bovine MTs | | | Yeast MTs | | |
| --- | --- | --- | --- | --- | --- | --- |
| | a (nm<sup>2</sup>/s) | $\alpha$ | b (nm<sup>2</sup>) | a (nm<sup>2</sup>/s) | $\alpha$ | b (nm<sup>2</sup>) |
| **Cin8Kin** | 2.03x10<sup>4</sup><br>±0.04x10<sup>4</sup> | **1.38**<br>**±0.01** | 5034<br>±550 | 1.16x10<sup>4</sup><br>±0.03x10<sup>4</sup> | **1.64**<br>**±0.01** | 1727<br>±435 |
| **Cin8+28NCKin** | 2.15x10<sup>4</sup><br>±0.04x10<sup>4</sup> | **1.29**<br>**±0.01** | 2278<br>±550 | 1.17x10<sup>4</sup><br>±0.03x10<sup>4</sup> | **1.29**<br>**±0.01** | 1844<br>±323 |

